# Mevalonate Biosynthesis is a Metabolic Vulnerability of Gemcitabine-resistant Pancreatic Cancer

**DOI:** 10.1101/2025.09.19.675739

**Authors:** Alica K. Beutel, Sabrina Calderon, Ethan Nghiem, Kevin Gulay, Leonor P. S. Santana, Eleni Zimmer, Rima Singh, Cecily Anaraki, Natalie Yousefian, Chantal Allgöwer, Gregory Tong, Cameron Geller, Alina Chao, Dennis Juarez, Thomas Seufferlein, Alexander Muir, Cholsoon Jang, Aimee L. Edinger, Marcus Seldin, Thomas F. Martinez, Alexander Kleger, David K. Chang, Herve Tiriac, Jennifer B. Valerin, David A. Fruman, Christopher J. Halbrook

**Affiliations:** Department of Molecular Biology and Biochemistry University of California Irvine, Irvine, USA; Department of Internal Medicine I, University Hospital Ulm, Ulm, Germany; Department of Surgery, University of California San Diego, La Jolla, USA; West of Scotland HPB Unit, Glasgow Royal Infirmary, Glasgow G61 1QH, UK; School of Cancer Sciences, University of Glasgow, Glasgow G61 1QH, UK; Institute of Molecular Oncology and Stem Cell Biology, University Hospital Ulm, Ulm, Germany; Department of Pharmaceutical Sciences University of California Irvine, Irvine, USA; Department of Developmental and Cell Biology University of California Irvine, Irvine, USA; Ben May Department for Cancer Research University of Chicago, Chicago, USA; Department of Biological Chemistry University of California Irvine, Irvine, USA; Chao Family Comprehensive Cancer Center, University of California Irvine, USA; Division of Interdisciplinary Pancreatology, Department of Internal Medicine I, University Hospital Ulm, Ulm, Germany; Department of Medicine, Division of Hematology and Oncology, University of California Irvine, Orange, USA

## Abstract

Pancreatic ductal adenocarcinoma (PDAC) is a lethal malignancy with a devastating prognosis. Gemcitabine, a pyrimidine anti-metabolite, is a cornerstone in PDAC therapy. However, resistance remains a major hurdle in clinical care. Resistance can arise from microenvironmental metabolites or through direct metabolic reprogramming of pancreatic cancer cells. Here, we generated PDAC models of acquired gemcitabine resistance to determine the relationship between these mechanisms. We observed that physiological levels of exogenous pyrimidines have a diminished ability to impact gemcitabine response in PDAC cells with acquired resistance. This occurs as the metabolic reprogramming of PDAC cells in response to gemcitabine treatment forces a suppression of the pyrimidine salvage pathway. Importantly, this metabolic rewiring renders gemcitabine-resistant PDAC cells highly susceptible to inhibition of the rate limiting enzyme of the mevalonate biosynthesis pathway, 3-hydroxy-3-methylglutaryl coenzyme A reductase (HMGCR), using statins. Notably, statin treatment inhibits the growth of gemcitabine-resistant tumors in immunocompetent mouse models. Through metabolite rescue experiments, we identified geranylgeranyl pyrophosphate as the critical metabolite lost during statin treatment, resulting in reduced protein geranylation in PDAC cells. Finally, as downregulation of the *HMGCR* is gradually acquired during gemcitabine resistance, we observed that *HMGCR* expression predicts patient response to gemcitabine. Collectively, these data demonstrate that the mevalonate biosynthesis pathway represents a promising therapeutic target in gemcitabine resistance and may serve as a biomarker to stratify treatment selection in PDAC patients.

## Introduction

Pancreatic ductal adenocarcinoma (PDAC) remains one of the deadliest major cancers (1). This is largely due to late diagnosis, aggressive tumor biology, scarcity of effective therapeutics, and early onset of resistance (2). Treatment of PDAC predominantly relies on conventional chemotherapy, with standard-of-care regimens leveraging the pyrimidine anti-metabolites gemcitabine and 5-fluorouracil, respectively. Chemotherapy combinations such as gemcitabine/nab-paclitaxel (3), FOLFIRINOX (5-fluorouracil, folinic acid, irinotecan, oxaliplatin) (4), or NALIRIFOX (5-fluorouracil, folinic acid, nanoliposomal irinotecan, oxaliplatin) (5) modestly extend survival and increase quality of life in advanced tumor stages, however, the lack of durable responses and rapid acquisition of resistance renders PDAC treatment particularly challenging.

The physiology of PDAC is partially responsible for its lack of therapeutic response. Pancreatic tumors are densely fibrotic and possess large populations of fibroblasts and immune cells (2). This creates high interstitial pressure that collapses functional vasculature, hindering diffusion of nutrients and chemotherapy (6–8). PDAC cells adapt to this nutritional challenge through metabolic strategies that conserve essential building blocks for cell proliferation and enhance nutrient scavenging (9, 10). As an example, we and others have shown that pancreatic cancer cells can obtain amino acids and glycans from their tumor microenvironment (11–15). We have also observed that PDAC cells avidly scavenge exogenous pyrimidines through nucleotide salvage pathways (16). This scavenging activity underlies the effectiveness of anti-pyrimidine therapies such as gemcitabine, a difluoro analog of deoxycytidine, widely used to treat PDAC and other cancers (17). Cancer cells efficiently import gemcitabine, which is incorporated into the DNA where it disrupts DNA synthesis through masked chain termination, evades DNA repair and induces apoptosis (17).

However, numerous mechanisms of gemcitabine resistance have been reported (17). Cell-intrinsic resistance mechanisms primarily involve metabolic reprogramming to reduce chemotherapy uptake, increase de novo pyrimidine synthesis, inhibit anti-metabolite chemotherapy by molecular competition, and enhance gemcitabine detoxification. As an example for non-cell-autonomous resistance that is driven by the tumor microenvironment, we and others have shown that nucleosides released by stromal and immune cells can directly impair the cytotoxic impact of gemcitabine on PDAC cells (16, 18). This occurs through molecular competition between exogenous deoxycytidine and gemcitabine at deoxycytidine kinase, a key step in the pyrimidine salvage pathway. Both cell-intrinsic and non-cell-autonomous resistance likely co-exist within pancreatic tumors during continuous gemcitabine treatment. Accordingly, it remains unknown if exogenous pyrimidines will retain a role in blocking the cytotoxicity of gemcitabine in the context of acquired resistance.

To examine this, we generated syngeneic murine PDAC cells with acquired gemcitabine resistance. Using these models, we found that physiological levels of deoxycytidine have reduced or no impact on the gemcitabine response of gemcitabine-resistant (GemR) PDAC cells. This results from extensive rewiring of multiple metabolic pathways during acquisition of chemoresistance that prevents nucleoside salvage. Interestingly, the adaptation to gemcitabine treatment downregulates the mevalonate biosynthesis pathway through suppression of SREBP2. This renders GemR PDAC susceptible to the statin family of small molecule inhibitors of 3-hydroxy-3-methylglutaryl coenzyme A (HMG-CoA) reductase (HMGCR), the rate-limiting enzyme in the mevalonate biosynthesis pathway. Finally, low expression levels of HMGCR in PDAC patients predicted poor response to gemcitabine treatment. Collectively, these data suggest that targeting the mevalonate biosynthesis pathway could be leveraged to improve treatment of PDAC.

## Results

### Physiological concentrations of exogenous deoxycytidine have diminished impact on gemcitabine response of resistant PDAC cells

To investigate the impact of gemcitabine resistance in PDAC within a system capable of interrogating both cell intrinsic and non-cell autonomous resistance mechanisms, we developed a model of acquired resistance using several murine PDAC cell lines derived from C57BL/6J *Kras*^+/LSL-G12D^; *Trp53*^+/R172H^; *Pdx1-Cre* (KPC) mice. Through a serial dose-escalation strategy (**Figure 1a**), we increased the gemcitabine tolerance of several KPC lines by several hundred-fold (**Figure 1b-e**; **Supplemental Fig. 1a,b**) generating cell lines that we termed ‘gemcitabine resistant’ (GemR). We next assessed if this resistance was mediated through a modulation of DNA damage or reduced cytotoxicity in response to gemcitabine treatment. Here, we observed that treatment with nanomolar concentrations of gemcitabine induced DNA damage, measured by γH2AX, and apoptosis, as indicated by caspase-3 cleavage, in parental but not GemR PDAC cells (**Figure 1f**). The gemcitabine resistance was maintained in vivo, as syngeneic GemR PDAC tumors treated with gemcitabine showed a marked loss of therapeutic response (**Figure 1g-i; Supplemental Fig. 1c-g**), and strikingly reduced levels of DNA damage compared to parental tumors (**Figure 1j**). Finally, consistent with our previous work (16), supplementation of parental PDAC cell lines with low micromolar concentrations of deoxycytidine shifted their gemcitabine sensitivity (**Figure 1k,l; Supplemental Fig. 1h**). However, the addition of physiological concentrations of deoxycytidine had markedly less impact on the dose response of GemR PDAC cell lines (**Figure 1k,l; Supplemental Fig. 1h**).

**Figure 1.**
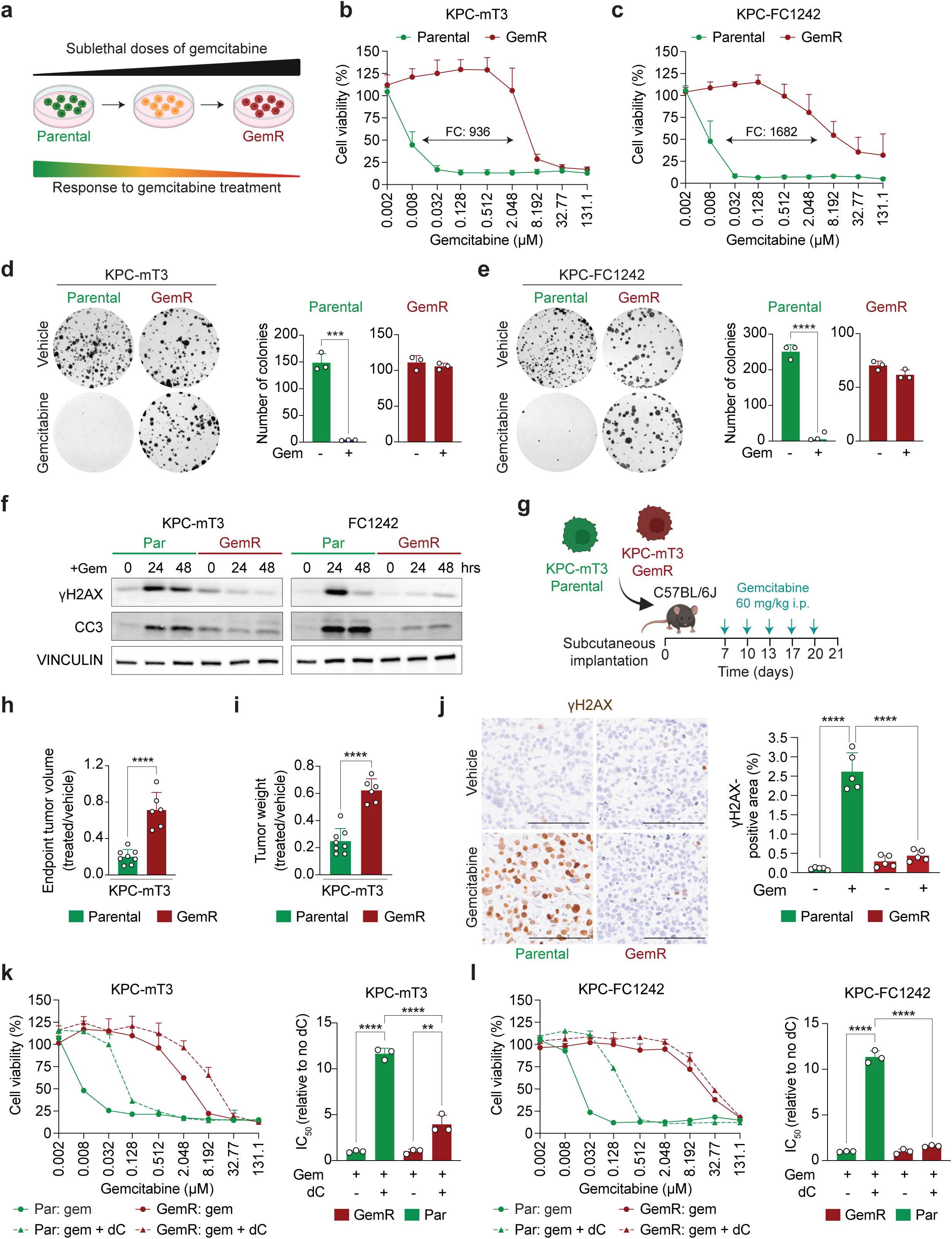
**a.** Schematic representation for a model of acquired gemcitabine resistance. **b-c**. Gemcitabine dose response curves for parental and gemcitabine-resistant (GemR) KPC-mT3 (**b**) and KPC-FC1242 (**c**) cells, with fold change (FC) indicated. **d-e.** Representative images and quantification of colony formation assays of vehicle or 0.01 µM gemcitabine (gem) treatment in parental and GemR KPC-mT3 (**d**), and KPC-FC1242 cells (**e**). **f.** Immunoblot of H2AX p-S139 (γH2AX) and cleaved Caspase 3 (CC3) on parental (Par) and GemR KPC-mT3 and KPC-FC1242 cells after 0, 24, and 48-hour treatment with vehicle or 100 nM gemcitabine (Gem). **g**. Treatment schematic of subcutaneous in vivo experiment with parental and GemR KPC-mT3 cells. **h-i**. Endpoint tumor volume (**h**) and tumor weight (**i**) comparison of parental and GemR KPC-mT3 allografts. **j**. Immunohistochemistry staining and quantification of γH2AX positive cells in resected tumors. **k-l.** Dose response curves and quantification for parental and GemR KPC-mT3 (**k**) and KPC-FC1242 (**l**) cells treated with gemcitabine +/- 10 µM deoxycytidine (dC). Error bars are presented as mean ± SD (**d, e, h, i, j, k, l**). *** *P* ≤ 0.001; **** *P* ≤ 0.0001 by two-tailed Student’s t-test (**d, e, h, i,**) and one-way ANOVA with Tukey post hoc test (**j, k, l**). Scale bar = 100 µm.

### Acquisition of gemcitabine resistance in PDAC cells downregulates the pyrimidine salvage pathway

The reduced impact of physiological pyrimidines on gemcitabine tolerance in GemR PDAC cells suggests either that the increased concentration of drug required to trigger a response may more effectively outcompete pyrimidines at the deoxycytidine kinase, or that the nucleoside salvage pathway, which is essential for gemcitabine activity, may have been reprogrammed. To investigate these possibilities, we performed a transcriptomic comparison between parental versus GemR PDAC cells. Principal component analysis revealed a clear separation between parental and GemR KPC-mT3 and KPC-1242 cells (**Figure 2a**). Interestingly GemR cells appeared to acquire gemcitabine tolerance through a combination of distinct and overlapping pathways. Consistent with this, we identified several hundred genes that were either upregulated or downregulated in resistant cells, with a subset of shared genes altered between the two cell lines (**Figure 2b**). Hierarchical clustering of the most significantly altered genes further distinguished parental from resistant cells (**Figure 2c**), confirming transcriptomic differences of gemcitabine sensitivity versus resistance and allowing us to assess these shared features across resistant states.

**Figure 2.**
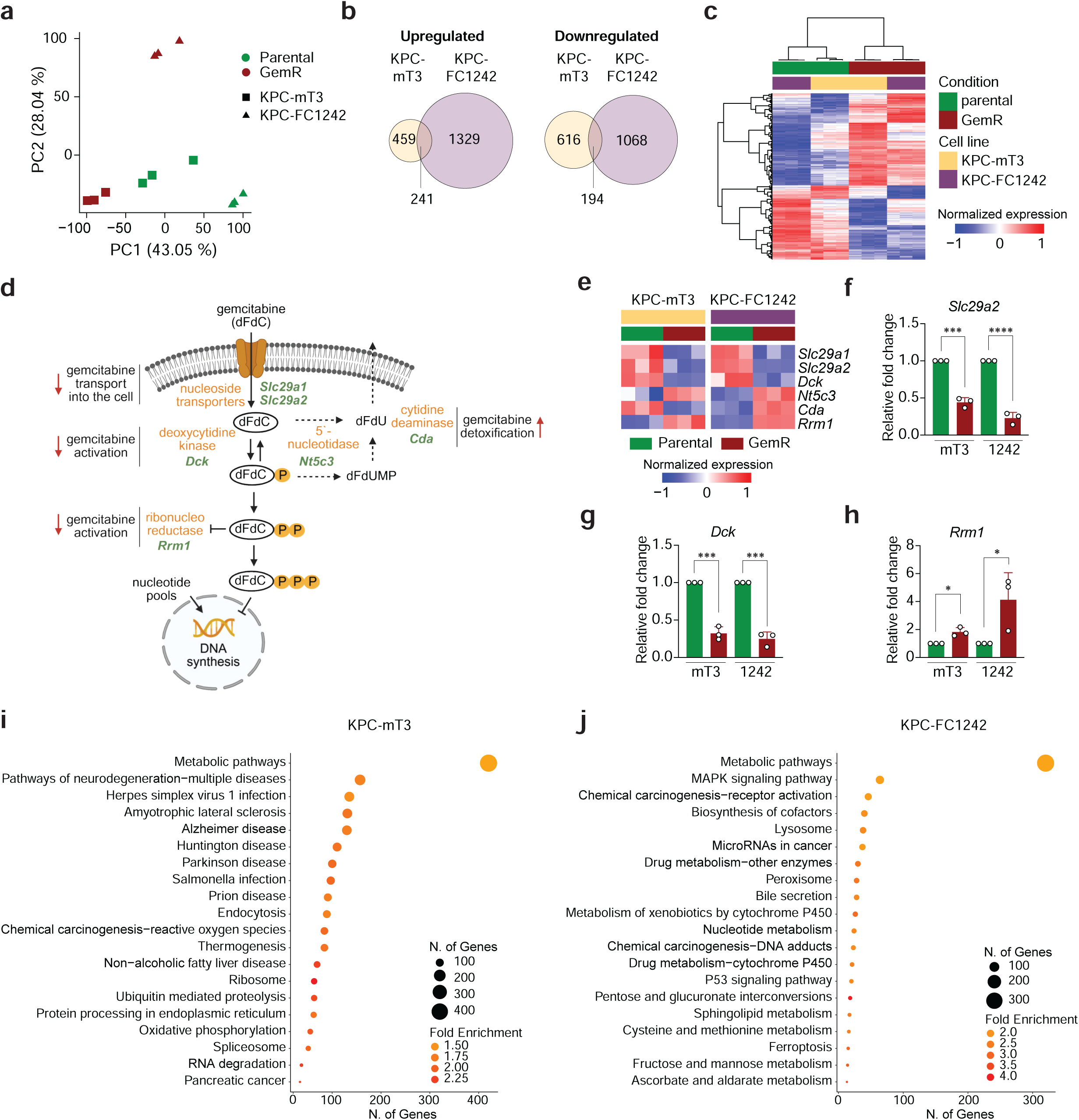
**a.** Principal component analysis of transcriptomic datasets of parental and gemcitabine-resistant (GemR) KPC-mT3 and KPC-FC1242 cells. **b**. Venn diagrams depicting the overlap of differentially expressed genes between parental and GemR KPC-mT3 and KPC-FC1242 cells. **c**. Heatmap displaying the intersect of differentially expressed genes in parental versus GemR KPC-mT3 and KPC-FC1242 cells. **d.** Schematic representation of gemcitabine metabolism. **e**. Heatmap displaying RNA-seq counts of selected genes involved in gemcitabine metabolism. **f-h**. qPCR validation of *Scl29a2* (**f**), *Dck* (**g**), and *Rrm1* (**h**) in parental versus GemR KPC-mT3 and KPC-FC1242 cells. **i-j.** KEGG pathway enrichment analysis of differentially expressed genes between parental and GemR KPC-mT3 (**i**) and KPC-FC1242 (**j**) cells, performed using ShinyGO. Error bars are presented as mean ± SD (**f, g, h**). * *P* ≤ 0.05; *** *P* ≤ 0.001; **** *P* ≤ 0.0001 by two-tailed Student’s t-test (**f-h**).

We first focused our analysis on genes associated with gemcitabine metabolism. Gemcitabine is a hydrophilic prodrug that requires transport into cells through specific nucleoside transporters, followed by sequential phosphorylation steps to exert cytotoxicity (**Figure 2d**). Our data indicate that GemR cells extensively rewire this pathway (**Figure 2e**). These changes include a downregulation of the equilibrative nucleoside transporters *Slc29a1* and *Slc29a2* in GemR PDAC cells, suggesting that the primary mechanism of resistance is through elimination of drug import at the cost of nucleotide salvage. The transcription of deoxycytidine kinase (*Dck*), required for pyrimidine salvage and gemcitabine activation, is also markedly reduced in GemR cells. In parallel, GemR cells upregulate the nucleotidase *Nt5c3* (in both KPC-mT3 and KPC-FC1242) and the deaminase *Cda* (in KPC-FC1242), both of which encode enzymes that inactivate and detoxify gemcitabine. Finally, to compensate for impaired pyrimidine salvage, GemR PDAC cells upregulate ribonucleotide reductase enzyme 1 (*Rrm1*), thereby increasing de novo nucleotide synthesis and reducing the incorporation of gemcitabine into the DNA. Targeted qPCR analysis validated the loss of nucleoside transport (**Figure 2f**), nucleotide capture and gemcitabine activation (**Figure 2g**), as well as increased de novo nucleotide synthesis (**Figure 2h**) in parental versus GemR PDAC cells.

Next, we performed KEGG pathway enrichment analysis on the differentially expressed genes of parental and GemR KPC-mT3 and KCP-1242 cells (**Figure 2i,j**). Concurrent with previous studies (19), we observed that metabolic pathways were among the most differentially expressed in these cells. Accordingly, we postulated that the metabolic rewiring accompanying gemcitabine resistance may represent an exploitable therapeutic vulnerability.

### Metabolic rewiring of GemR PDAC suppresses the mevalonate biosynthesis pathway

Given the importance of metabolic pathways in our differential gene expression analysis, we performed a targeted inhibitor screen to identify potential actionable vulnerabilities (**Figure 3a,b**). Here, we found that several compounds exhibited greater efficacy in GemR compared to parental PDAC cells. Consistent with previous studies, we observed that GemR KPC-mT3 cells were more sensitive to 2-deoxy-D-glucose (2-DG) inhibition of glycolysis and dihydroorotate dehydrogenase (DHODH) inhibition of de novo pyrimidine synthesis via BAY-2402234 (19). Conversely, GemR KPC-mT3 cells appeared less sensitive to BPTES and CB-839, both of which target glutamine metabolism, a pathway previously described as a central hallmark of the metabolic reprogramming of PDAC cells driven by oncogenic KRAS (20). Interestingly, GemR KPC-mT3 cells displayed increased sensitivity to the HMG-CoA reductase (HMGCR) inhibitor pitavastatin (**Figure 3a**). Similarly, GemR KPC-FC1242 cells were more susceptible not only to pitavastatin but also to the sterol regulatory element-binding protein 2 (SREBP2) inhibitor fatostatin, whereas additional metabolic compounds tested in KPC-FC1242 cells did not reveal further vulnerabilities (**Figure 3b**).

**Figure 3.**
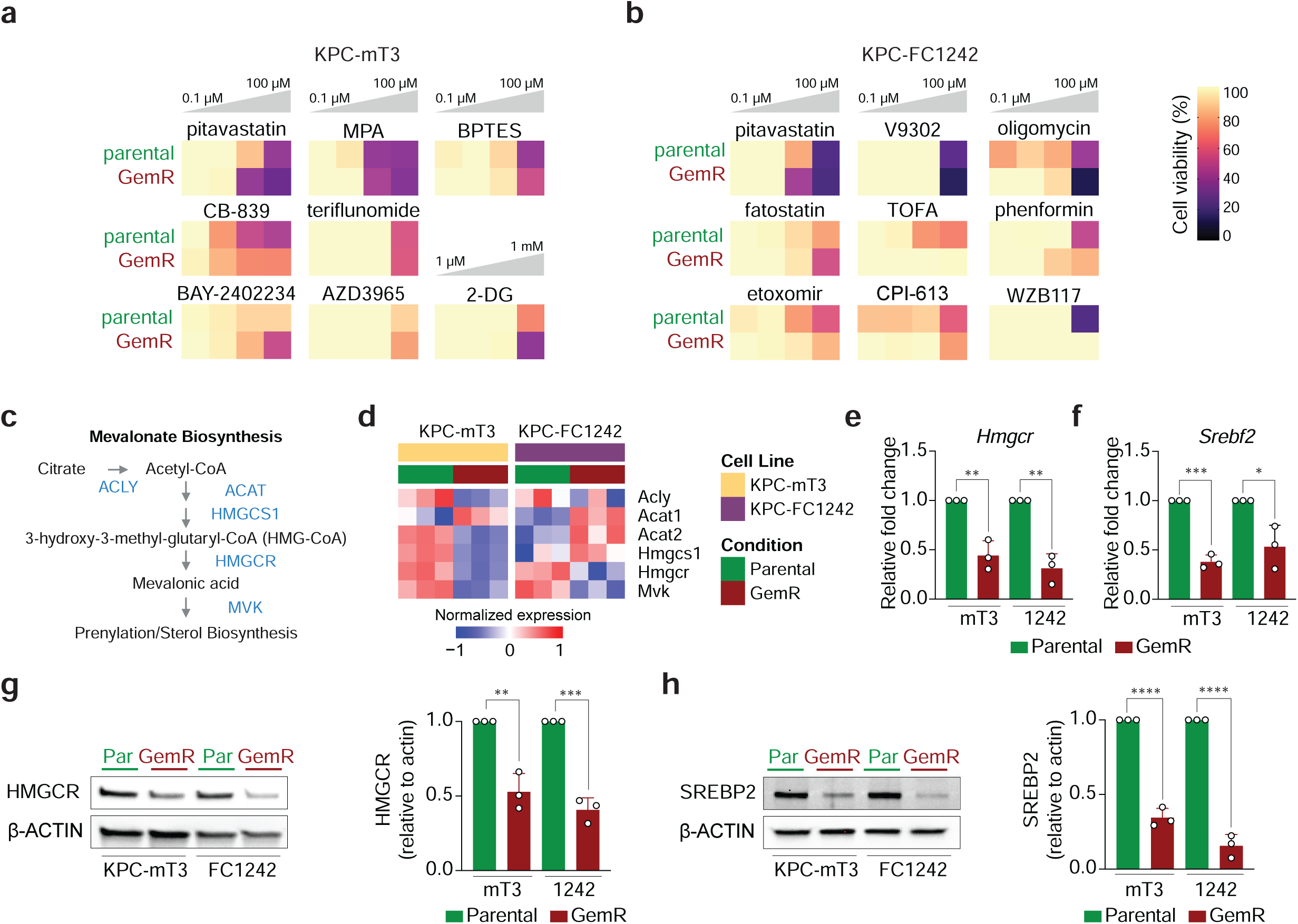
**a-b.** Customized drug viability assay screen on parental versus gemcitabine-resistant (GemR) KPC-mT3 (**a**) and KPC-FC1242 (**b**) cells with increasing concentrations (0.1 µM - 100 µM) of metabolic inhibitors. **c**. Simplified overview of the mevalonate pathway. **d.** Heatmap displaying RNA-seq counts of selected genes involved in the mevalonate pathway in parental versus GemR KPC-mT3 and KPC-FC1242 cells. **e-f.** qPCR validation of *Hmgcr* (**e**) and *Srebf2* (**f**) in parental versus GemR KPC-mT3 and KPC-FC1242 cells. **g.** Representative immunoblot for HMGCR and quantification of relative HMGCR levels normalized to β-ACTIN levels on parental and GemR KPC-mT3 and KPC-FC1242 cell lysates. **h.** Representative immunoblot for SREBP2 and quantification of relative SREBP2 levels normalized to β-ACTIN levels on parental and GemR KPC-mT3 and KPC-FC1242 cell lysates. Error bars are presented as mean ± SD (**e, f, g, h**). * *P* ≤ 0.05; ** *P* ≤ 0.01; *** *P* ≤ 0.001; **** *P* ≤ 0.0001 by two-tailed Student’s t-test (**e, f, g, h**). MPA, mycophenolic acid; 2-DG, 2-deoxy-D-glucose, TOFA, Tofacitinib.

We chose to focus on the pitavastatin hit for multiple reasons. First, gene set enrichment analysis in GemR KPC-mT3 cells showed that cholesterol biosynthesis was more enriched in parental than in resistant cells (**Supplementary Figure 2a**), indicating a relevant rewiring of the mevalonate biosynthesis pathway during resistance. HMGCR, the target of the statins, is widely used for primary and secondary prevention of cardiovascular disease. Beyond their lipid-lowering role, statins possess several attractive features as anti-cancer therapeutics, with potential to block the biosynthesis of many critical metabolites (21). Furthermore, dependence on the mevalonate biosynthesis pathway has emerged as a drug resistance mechanism in several cancer models (22–24), and combining statins with standard-of-care treatment has shown promising results in early clinical trials in blood cancers (25) and PDAC (26).

HMGCR is the rate-limiting enzyme of the mevalonate biosynthesis pathway that produces metabolic precursors for sterol synthesis and prenylation moieties required for membrane anchorage of proteins (**Figure 3c**). Assessing our transcriptomic data revealed that several enzymes in this pathway were downregulated in GemR PDAC cells (**Figure 3d**). The mevalonate biosynthesis pathway is largely regulated by SREBP2 (encoded by the *Srebf2* gene). Interestingly, both *Hmgcr* and *Srebf2* expressions were markedly reduced in GemR PDAC cells (**Figure 3e,f)**. Immunoblotting confirmed that these transcriptomic changes translated to lower levels of both HMGCR and SREBP2 total protein in GemR as compared to parental PDAC cell lines (**Figure 3g,h; Supplementary Figure 2b**).

### PDAC acquires sensitivity to mevalonate biosynthesis pathway inhibition while gaining gemcitabine tolerance

Our initial inhibitor screen relied on ATP levels or mitochondrial enzyme activity as a surrogate for cell number. To validate these findings, we directly tested whether downregulation of HMGCR expression sensitizes PDAC cells to statin treatment, which has been reported in other cancer models (23). Indeed, GemR PDAC cells were highly vulnerable to pitavastatin, exhibiting greater toxicity compared to their parental counterparts as determined by colony formation assays (**Figure 4a,b**). Furthermore, GemR PDAC cells were far more sensitive to a panel of statins, including pitavastatin, atorvastatin, simvastatin and fluvastatin (**Figure 4c; Supplementary Figure 3a-c**), confirming a broad vulnerability to this class of inhibitors. Among these compounds, we selected pitavastatin for subsequent experiments due to its superior pharmacokinetic properties for targeting cancer (27). Pitavastatin is a lipophilic statin with a relatively long half-life and limited metabolism via the cytochrome P450 system, which reduces potential drug-drug interactions (28). Interestingly, we noted that the degree of gemcitabine resistance gained by PDAC cells during serial dosing was inversely correlated to their sensitivity to statins, further demonstrating the interplay between these two pathways (**Figure 4d-h; Supplementary Figure 4a-c**).

**Figure 4.**
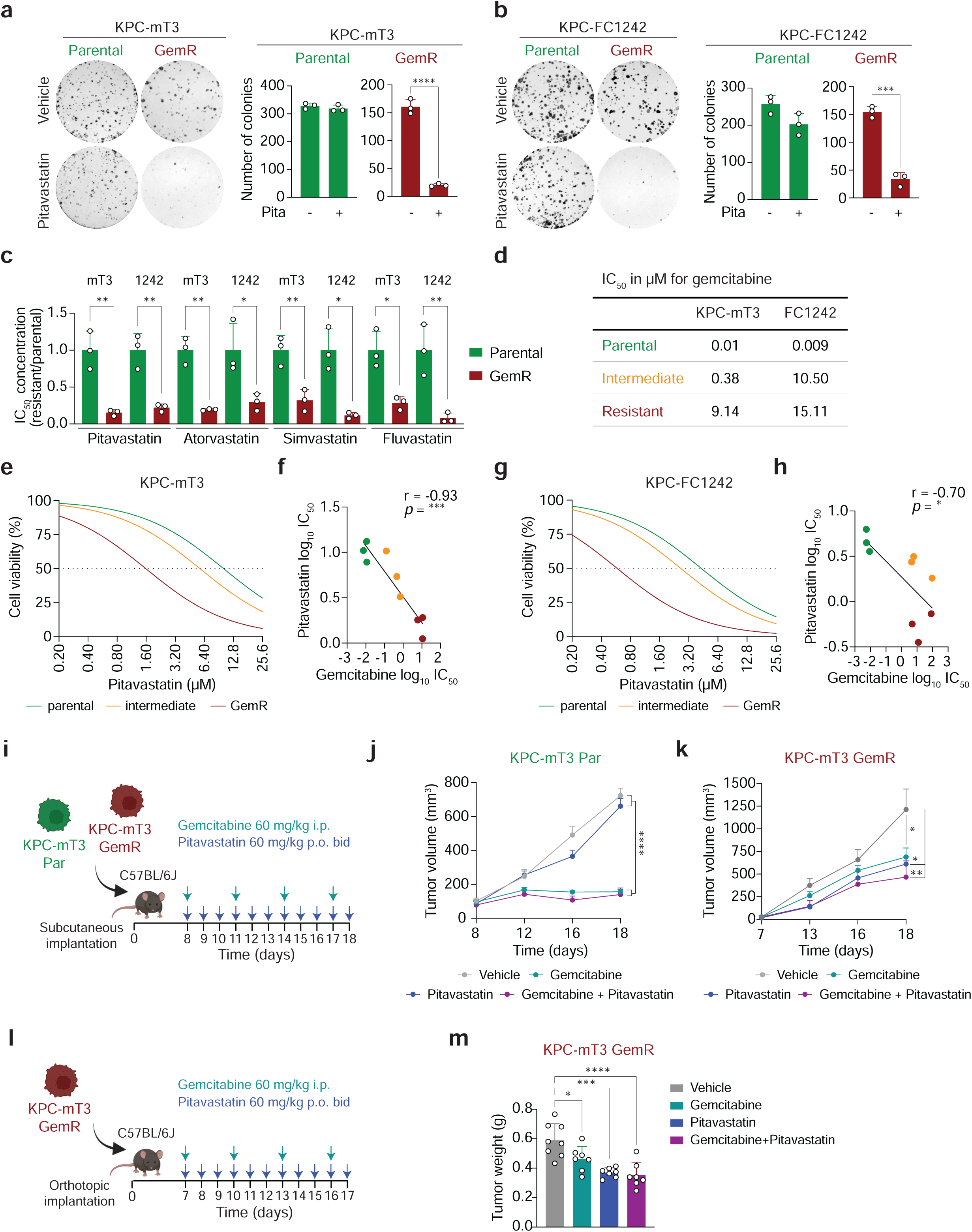
**a-b.** Representative images and quantification of colony formation assays of vehicle or 0.8 µM pitavastatin (pita) treatment in parental and gemcitabine-resistant (GemR) KPC-mT3 (**a**) and KPC-FC1242 (**b**) cells. **c.** IC_50_ concentrations for pitavastatin, atorvastatin, simvastatin, and fluvastatin normalized to gemcitabine parental KPC-mT3 and KPC-FC1242 cells. **d.** Table with IC_50_ concentrations for gemcitabine in parental, intermediate resistant, and GemR KPC-mT3 and KPC-FC1242 cells. **e-f**. IC_50_ curves for pitavastatin (**e**) and correlation of IC_50_ concentrations for pitavastatin and gemcitabine (**f**) of parental, intermediate resistant, and GemR KPC-mT3 cells. **g-h**. IC_50_ curves for pitavastatin (**g**) and correlation of IC_50_ concentrations for pitavastatin and gemcitabine (**h**) in parental, intermediate resistant and GemR KPC-FC1242 cells. **i.** Treatment schematic for subcutaneous parental and GemR KPC-mT3 tumors in C57BL/6J mice. **j-k**. Tumor growth kinetics of parental (**j**) and GemR (**k**) KPC-mT3 allografts in response to vehicle, gemcitabine (60 mg/kg i.p. every three days), pitavastatin (60 mg/kg p.o. twice daily), or combined gemcitabine and pitavastatin treatment. **l.** Treatment schematic for orthotopic GemR KPC-mT3 tumors in C57BL/6J mice. **m.** Final tumor mass of orthotopic KPC-mT3 allografts after treatment with vehicle, gemcitabine (60 mg/kg i.p. every three days), pitavastatin (60 mg/kg p.o. twice daily), or combined gemcitabine and pitavastatin treatment. Error bars are presented as mean ± SD (**b, c, m**) or mean ± SEM (**j, k**). * *P* ≤ 0.05; ** *P* ≤ 0.01; *** *P* ≤ 0.001; **** *P* ≤ 0.0001 by two-tailed Student’s t-test (**b, c**), Pearson correlation (**f, h**), and one-way ANOVA with Tukey post hoc test (**j, k, m**).

Assessing response to statins in mice is complicated by the fact that standard rodent diet contains high levels of geranylgeraniol that rescues statin-induced anticancer activity (29). Using a liquid diet that precludes geranylgeraniol (**Figure 4i**), we observed that parental KPC-mT3 allografts failed to exhibit any therapeutic response to pitavastatin treatment (**Figure 4j, Supplementary Figure 4d**). In contrast, GemR PDAC tumors responded to both single-agent and combination treatments with pitavastatin (**Figure 4k, Supplementary Figure 4e**). Importantly, this response was maintained in orthotopic GemR KPC-mT3 tumors (**Figure 4l,m**). The extent of mouse body weight loss remained within acceptable limits for single-agent and combination treatments (**Supplementary Figure 4f**), suggesting in vivo tolerability of these therapies.

### Maintenance of geranylgeranyl pyrophosphate pools is critical for PDAC

The mevalonate biosynthesis pathway has numerous critical outputs required for cell survival (**Figure 5a**) (21). These include cholesterol, squalene, co-enzyme Q10 (CoQ10), farnesyl pyrophosphate (FPP), geranylgeranyl pyrophosphate (GGPP), among others. Both FPP and GGPP are used for post-translational prenylation, a modification necessary for membrane anchoring of many small GTPases. FPP primarily mediates membrane anchoring of RAS proteins, including oncogenic KRAS that is central to PDAC pathology, whereas GGPP is more associated with RHO-family proteins (21).

**Figure 5.**
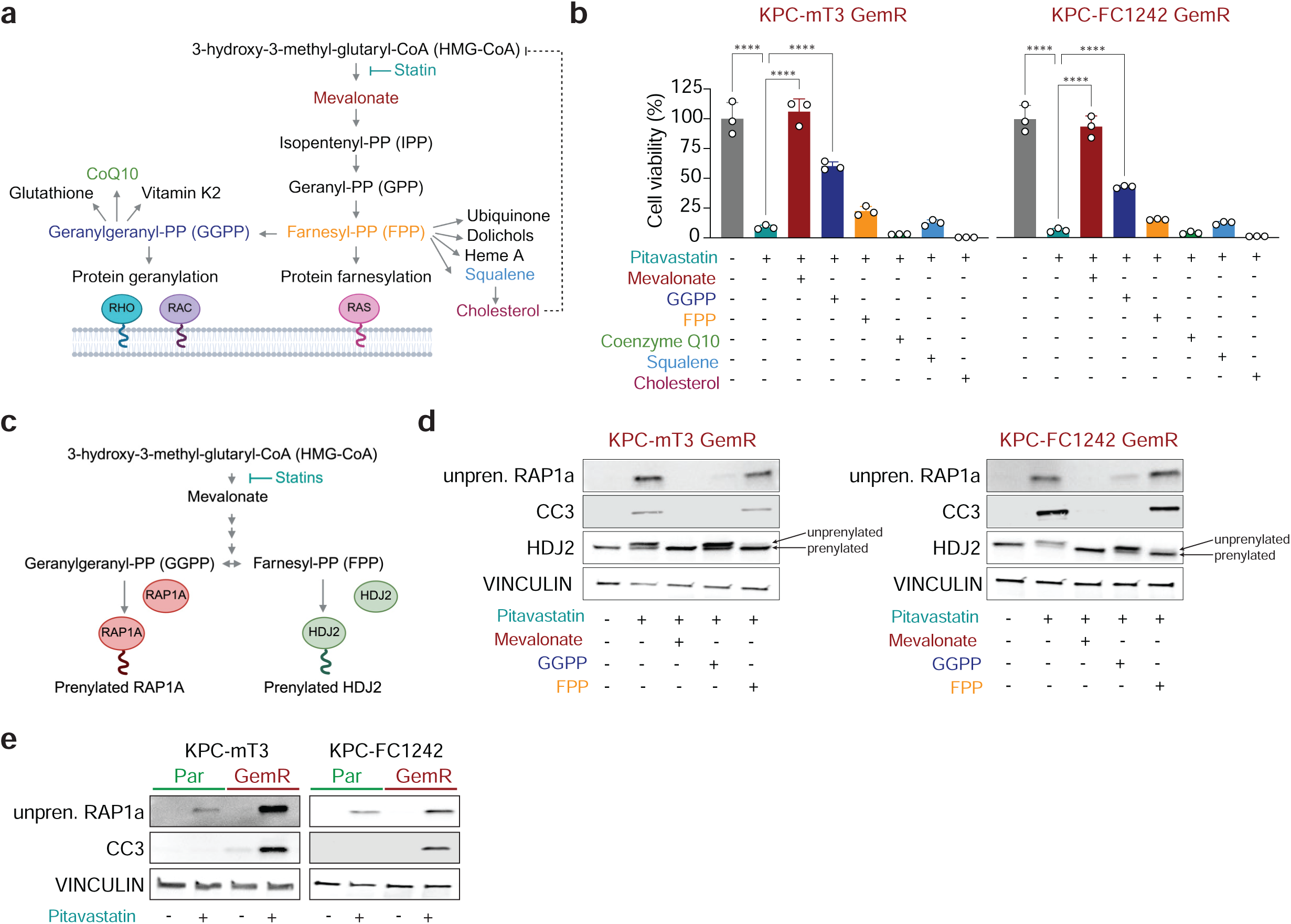
**a.** Schematic representation of the mevalonate pathway. **b.** Cell viabilities of gemcitabine-resistant (GemR) KPC-mT3 and GemR KPC-FC1242 cells treated with 10 µM pitavastatin alone or in combination with 1 mM mevalonate, 20 µM GGPP, 20 µM FPP, 10 µM coenzyme Q10, 50 µM squalene, or 50 µM cholesterol. Compounds used for rescue experiments are indicated by color scheme in (Figure 5a). **c.** Schematic representation of prenylation of RAP1a and HDJ2. **d.** Immunoblot for unprenylated RAP1a, Cleaved Caspase 3 (CC3) and HJD2 in parental and GemR KPC-mT3 or KPC-FC1242 cells after 24-hour treatment with 5 µM pitavastatin alone or in combination of 1 mM mevalonate, 20 µM GGPP, or 20 µM FFP. **e.** Immunoblot comparison of unprenylated RAP1a and CC3 levels in parental (Par) and GemR KPC-mT3 and KPC-FC1242 cells in response to 5 µM treatment with pitavastatin. Error bars are presented as mean ± SD (**b**). **** *P* ≤ 0.0001 by one-way ANOVA with Tukey post hoc test (**b**).

To determine the critical downstream nodes of the mevalonate biosynthesis pathway affected by statin treatment in PDAC, we performed metabolite addback experiments (**Figure 5b, Supplementary Figure 5a**). Here, we observed that direct downstream supplementation of mevalonate nearly completely rescued GemR PDAC cells from pitavastatin-induced cytotoxicity. In contrast, downstream metabolites like FPP, CoQ10, and squalene failed to restore GemR PDAC growth when challenged with pitavastatin. Cholesterol further increased the cytotoxicity of pitavastatin, likely due to its negative feedback inhibition on HMGCR (30) (**Supplementary Figure 5b)**. In contrast, GGPP robustly rescued GemR PDAC cells from statin treatment, highlighting GGPP as a critical metabolic node of PDAC survival.

To confirm that our metabolite additions restored their intended pools during these rescue experiments, we employed our previous approach of assessing the prenylation of FPP- or GGPP-modified proteins (22). Here, we used an antibody specific for unprenylated RAP1a, a canonical GGPP-modified protein, as well as an antibody that recognizes both prenylated and unprenylated HDJ2, which produces a sufficiently large gel shift to visualize both forms **(Figure 5c)**. Treatment with pitavastatin led to accumulation of unprenylated RAP1A and HDJ2, accompanied by caspase-3 cleavage in GemR PDAC cells (**Figure 5d**). The addition of mevalonate fully restored prenylation of both pools, whereas GGPP and FPP addition specifically restored prenylation pools of RAP1A or HDJ2, respectively. Importantly, only mevalonate and GGPP addition prevented caspase-3 cleavage, mirroring the rescue of cell growth observed in **Figure 5b**. Finally, we found that treatment of parental versus GemR PDAC cells with the same concentration of pitavastatin resulted in a more pronounced loss prenylation of RAP1a and selective induction of caspase-3 cleavage in the resistant cells (**Figure 5e**).

### HMGCR expression correlates with gemcitabine response in PDAC patients

To further explore the relationship of the mevalonate biosynthesis pathway and gemcitabine response in patients, we analyzed the International Cancer Genome Consortium (ICGC) dataset, that includes patients who underwent primary surgical resection for PDAC and received gemcitabine-based adjuvant chemotherapy (31). Interestingly, gemcitabine uptake and metabolism genes did not impact overall survival (OS) in PDAC patients. Gemcitabine treatment conferred an OS benefit in patients irrespective of whether gemcitabine uptake expression levels were high or low (**Figure 6a,c**). *HMGCR* expression levels were also not prognostic (**Figure 6b,d**). However, in patients with low *HMGCR* expression, treatment with gemcitabine provided no survival benefit and was even associated with worse outcomes, consistent with our experimental model of gemcitabine resistance (**Figure 6b,d**). In contrast, patients with high *HMGCR* gene expression, predicted by our models to be gemcitabine-sensitive, showed a marked improvement in OS with gemcitabine-based treatment (**Figure 6b,d**).

**Figure 6.**
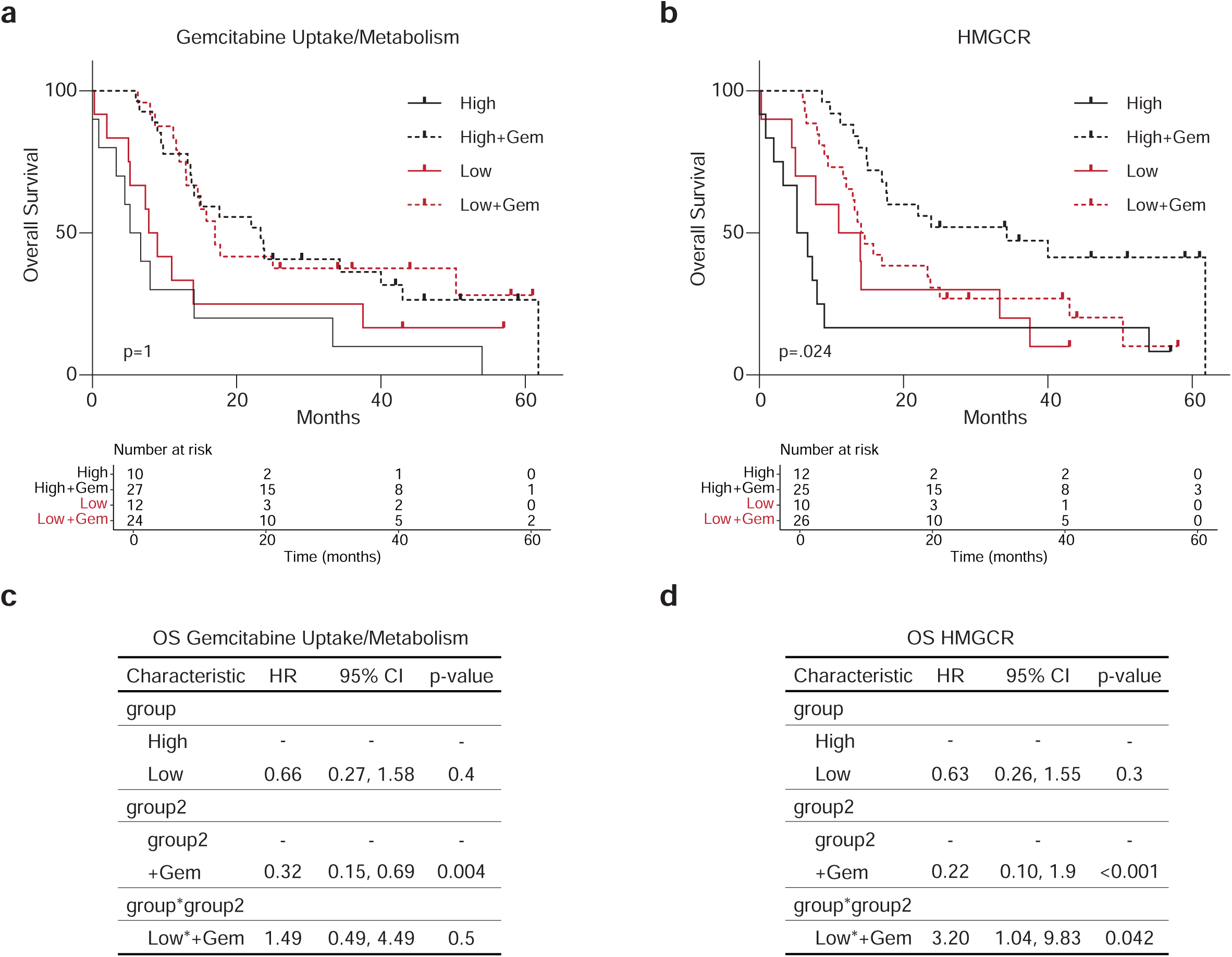
**a.** Kaplan-Meier curves of overall survival of untreated pancreatic cancer patients with either high (n=10) or low gemcitabine uptake/activation gene signature expression (n=12), compared to gemcitabine treated patients (high n=27, low n=24). **b.** Kaplan-Meier curves of overall survival of untreated pancreatic cancer patients with either high (n=12) or low *HMGCR* gene expression (n=10), compared to gemcitabine treated patients (high n=25, low n=26). **c-d.** Table depicting results from the cox proportional hazard model of gemcitabine uptake/metabolism (**c**) and *HMCGR* expression (**d**) on overall survival (OS) with or without gemcitabine treatment. HR, hazard ratio; CI, confidence interval.

Our experimental data indicate that the acquisition of gemcitabine resistance in PDAC correlates with a downregulation of *HMGCR* expression. This finding is mirrored in the ICGC cohort, where low *HMGCR* expression, a surrogate marker for gemcitabine resistance, predicts poor response to gemcitabine-based chemotherapy, highlighting its potential as a predictive biomarker.

## Discussion

Our findings indicate that the mevalonate biosynthesis pathway is a key metabolic node in pancreatic cancer that is rewired in acquired gemcitabine resistance. Blockade of this pathway has been shown to enhance the efficacy of the BCL-2 inhibitor venetoclax in leukemia (22), sensitize triple-negative breast cancer to AKT inhibitors (23), and overcome resistance to etoposide and cisplatin in small cell lung cancer (24). Interestingly, despite the production of numerous important metabolites, our studies indicate that the key output is the production of GGPP. In fact, the enzyme responsible for GGPP production, GGPS1, has previously emerged from CRISPR screens as a potential metabolic vulnerability in PDAC (32). However, specific targeting of the GGPP modification of proteins through geranylgeranyltransferase (GGTase) inhibitors has lagged in development as compared to the efforts to develop farnesyltransferase inhibitors (FTPase), and dual FTPase and GGTase inhibitors have demonstrated poor tolerability (33). Accordingly, upstream inhibition of HMGCR using statins, that are widely prescribed and generally well tolerated, represents a more practical therapeutic strategy. Additionally, approaches such as vertical targeting of the mevalonate biosynthesis pathway using compounds like bisphosphonates (34) could be explored to further enhance the potency of statins in PDAC treatment.

Preclinical evidence supports statins as anti-cancer therapeutics, with the exception of one murine study linking long-term statin treatment to EMT (35). However, clinical studies in PDAC patients have produced conflicting results (21). Although observational studies and meta-analyses reported increased survival in PDAC patients using statins (36–39), a randomized phase 2 trial found that adding low-dose simvastatin to gemcitabine provided no clinical benefit in patients with advanced disease (40). This lack of efficiency may reflect insufficient intratumoral target engagement rather than pathway irrelevance, properties that can be influenced by statin choice, statin dose and dietary factors, all of which are critical for effective pathway inhibition (27). Pitavastatin, in particular, exhibits stronger mevalonate pathway inhibition in cancer models and pharmacokinetic properties that support improved tumor exposure (28). Moreover, exogenous isoprenoids can restore protein geranylgeranylation and rescue statin-induced cytotoxicity. For example, geranylgeraniol is found in white rice and standard murine chows (29, 41), compounding the interpretation of both clinical and preclinical statin treatments. Careful consideration of statin type, dosing and dietary factors may help translate preclinical benefits into clinical outcomes. Indeed, early-phase studies such as combination of pitavastatin and venetoclax in blood cancer have shown promise (25). A recent single-arm, phase 2 clinical trial (NCT06241352) evaluated the addition of atorvastatin to standard-of-care chemotherapy in 42 advanced PDAC patients whose tumor markers plateaued as an indication of resistance. The addition of atorvastatin to the existing chemotherapy regimen resulted in significant reductions in tumor markers in 70% of patients suggesting improved chemotherapy efficacy (26). Additional trials are currently investigating the use of statins (NCT04245644) or a combination of cholesterol-disrupting agents (NCT04862260) in PDAC, highlighting ongoing clinical interest in targeting the mevalonate pathway.

Interestingly, beyond the impact of statins on patient survival, several meta-analyses reported lower occurrence of PDAC associated with statin use (36, 42, 43). While our data and that of others suggest that the primary downstream metabolic pathway impacted by statin treatment in the context of drug resistance is GGPP production and reduced protein geranylation, as opposed to cholesterol production reported in one PDAC study (26), other outputs of the mevalonate biosynthesis pathway affected by HMGCR blockade may also influence pancreatic tumorigenesis. For example, amplification of RAS signaling has been shown to play an essential role in pancreatic intraepithelial neoplasia (PanIN) formation and progression to PDAC (44, 45). Here, chronic deprivation of FPP and GGPP through statin treatment could limit KRAS prenylation and thus impair the initiation of feed-forward oncogenic signaling. Additionally, statins have been associated with suppression of chronic pancreatic inflammation, which is required for tumorigenesis in murine models (46, 47). Accordingly, statins offer potential avenues to prevent development or progression of PDAC by acting on the precancerous epithelium and microenvironment.

There are multiple opportunities to repurpose statins in the treatment of pancreatic cancer. Apart from a potential cancer prevention strategy, combining pitavastatin and gemcitabine-based treatment has potential to leverage the combinational effect and potentially delay any onset of gemcitabine resistance. We will soon evaluate this in a single-arm Phase 1b dose-finding study using a Bayesian Optimal Interval design, treating unresectable PDAC patients with pitavastatin in combination with gemcitabine and nab-paclitaxel. Alternatively, pitavastatin can be combined with second-line chemotherapy regimens after progression on gemcitabine-based treatment, exploiting the increased sensitivity to mevalonate biosynthesis pathway inhibition in resistance.

Despite tremendous efforts to use transcriptomic data to stratify PDAC patients into subtypes that could be used to inform treatment (2), no reliable biomarkers exist in routine clinical practice. Consequently, treatment of advanced PDAC still largely relies on empiric chemotherapy combinations. The choice of regimen is primarily determined by patient performance status and comorbidities, often resulting in limited efficacy and substantial toxicity. Although precision oncology approaches leveraging molecular, cellular, or functional tumor analyses are emerging to optimize therapeutic efficacy and minimize toxicity, validated biomarkers to guide clinical decision-making are still lacking. Our retrospective analysis of sequencing data revealed that *HMGCR* gene expression has potential to serve as a predictive biomarker for patient response to gemcitabine-based chemotherapy. Low *HMGCR* expression could identify patients unlikely to benefit from gemcitabine-based treatments, who may respond better to 5-fluorouracil/oxaliplatin/irinotecan-based regimens and to statins. Beyond gene expression, HMGCR levels could be assessed through patient tumor tissue staining or liquid biopsy, enabling dynamic monitoring of resistance before tumor progression becomes evident on imaging. Thus, standardized assessment of *HMGCR* expression levels could inform treatment selection and identify patients susceptible to statin therapy.

In summary, our findings highlight the therapeutic relevance of mevalonate pathway dependency in PDAC and support the potential for personalized treatment strategies that warrant further clinical evaluation.

## Materials and Methods

### Study approvals

All animal procedures were performed at the University of California, Irvine in compliance with the Institution Animal Care and Use Committee (AUP-23-084) and the Institutional Biosafety Committees (BUA-R315). Male C57BL/6J mice were bred in house at the University of California Irvine and purchased from Jackson Laboratory. Mice were housed and bred in conventional health status-controlled animal facilities. All mouse work aspects were carried out following strict guidelines to ensure careful, standardized, consistent, and ethical handling of mice.

### Cell culture

Pancreatic cancer lines were established from C57BL/6J Kras^+/LSL-G12D^; Trp53^+/LSL-R172H^; Pdx-1-Cre (KPC) mice. The designated KPC-mT3, KPC-FC1242, and KPC-FC1199 cell lines were obtained from Dr. David Tuveson. Cell lines were cultured in DMEM containing 10% FBS and propagated at 37°C under 5% (v/v) CO_2_ atmosphere.

### Generation of gemcitabine resistance

To generate an acquired gemcitabine resistance, KPC-mT3, KPC-FC1242 and KPC-FC1199 cells were treated twice per week with increasing concentrations of sublethal doses of gemcitabine (Selleckchem) for several months. Cell viability and half maximal inhibitory concentrations (IC_50_) were assessed regularly until a desired fold change was obtained.

### Cell viability assay

KPC-mT3, KPC-FC1242, or KPC-FC1199 cells were seeded at 2,000 cells per well in triplicates into white bottom 96-well plates (Corning). Twenty-four hours after seeding, cells were treated in triplicate with cytotoxic agents for 72 hours. Cell viability was analyzed using Cell Titer Glo 2.0 Assay (Promega) according to the manufacturer’s protocol. Luminescence was measured for 1000 ms using a microplate reader (Varioskan LUX™, Thermo Scientific). Viability percentages were normalized to vehicle-treated cell viability. Cell viability curves were generated using GraphPad Prism (GraphPad Software). Cell viability for IC_50_ determination for gemcitabine and statins was assessed in triplicates across three independent experiments. Gemcitabine, pitavastatin, simvastatin, atorvastatin, and fluvastatin were purchased from Selleckchem.

### Colony formation assay

KPC-mT3, KPC-FC1242, or KPC-FC1199 cells were seeded at 1,000 cells per well in triplicates into 6-well plates (Corning) and treated 24 hours after seeding with gemcitabine (0.01 µM) or pitavastatin (0.8 µM). Following cultivation for seven to ten days, cells were fixed with methanol and stained with 1% crystal violet solution (Sigma-Aldrich). Number of colonies (defined as > 0.1 mm^2^) were quantified using ImageJ software (National Institutes of Health).

### Rescue experiment

KPC-mT3, KPC-FC1242, or KPC-FC1199 gemcitabine-resistant cells were seeded at 2,000 cells per well in triplicates into white bottom 96-well plates (Corning) for 24 hours. Cells were treated with either 3 µM, 5 µM or 10 µM of pitavastatin and supplemented with mevalonic acid (1 mM), GGPP (20 µM), FPP (20 µM), coenzyme Q10 (10 µM), squalene (50 µM), or cholesterol (50 µM) for 72 hours. Cell viability was analyzed using Cell Titer Glo 2.0 Assay (Promega) according to the manufacturer’s protocol. Luminescence was measured for 1000 ms using a microplate reader (Varioskan LUX™, Thermo Scientific). Viability percentages were normalized to vehicle-treated cell viability. Cell viability graphs were generated using GraphPad Prism (GraphPad Software). Mevalonic acid, squalene and cholesterol were purchased from Sigma; coenzyme Q10, GGPP, and FPP were purchased from Cayman Chemical.

### Metabolic inhibitor screen

KPC-mT3 or KPC-FC1242 gemcitabine-resistant cells were seeded at 2,000 cells per well in duplicates into 96-well plates for 24 hours. Cells were treated with the indicated metabolic inhibitors at a concentration range of 0.1 µM to 100 µM for 72 hours. Cell viability was analyzed using Cell Titer Glo 2.0 Assay (Promega) for KPC-mT3 or MTT assay (Sigma-Aldrich) for KPC-FC1242 cells according to the manufacturer’s protocol. Luminescence was measured for 1000 ms using a microplate reader (Varioskan LUX™, Thermo Scientific) for Cell Titer Glo Assay, and absorbance was measured at 590 nm wavelength using a spectrophotometer (Tecan Infinite M200 Pro) for MTT assay, respectively. Viability percentages were normalized to vehicle-treated cell viability.

### Isolation of RNA

RNA was collected from parental and gemcitabine-resistant cells in a 6-well plate in triplicates using the RNeasy Plus Mini Kit (Qiagen). RNA yield was quantified via the NanoDrop™ 2000/2000c Spectrophotometers (Thermo Fisher).

### RNA sequencing

Samples were normalized and submitted to the UCI Genomics High Throughput Facility at University of California, Irvine for bulk RNA sequencing. RNA quality was confirmed on an Agilent Bioanalyzer to have RIN quality > 9. Stranded cDNA libraries were sequenced to a read depth of 40 million 150bp reads per sample on an Illumina NovaSeq 6000 on an S4 chip.

### Transcriptomic analysis

FastQFiles were aligned using STAR, bedgraphs were generated, and RNA counts were obtained in raw count format. RNA-seq data were analyzed using the *limma* framework (48). Raw counts were normalized using the Trimmed Mean of M-values (TMM) method, transformed to log_2_ counts per million (log_2_CPM), and mean-variance relationships were estimated with *voom* transformation. Lowly expressed genes were filtered via k-means clustering (*k* = 2) on mean log_2_CPM values. Principal component analysis (PCA) was performed on the entire feature space. Differential expression between parental and gemcitabine-resistant conditions was assessed using linear modeling with empirical Bayes variance moderation. Significance was determined at an adjusted p-value < 0.05 (Bonferroni correction). Gene set enrichment analysis (GSEA) was performed with the *ClusterProfiler* package (49) using Gene Ontology (GO) and Reactome pathway databases, with multiple testing correction by the Benjamini-Hochberg method.

KEGG pathway enrichment analysis on differentially expressed genes of parental and gemcitabine-resistant KPC-mT3 and KPC-FC1242 cells was performed using ShinyGO (50).

### Patient survival data

Patients were identified from the Australian Pancreatic Genome Initiative’s (APGI) contribution to the International Cancer Genome Consortium’s (ICGC) Pancreatic Cancer project (51) and the corresponding RNAseq data was downloaded from the last ICGC DCC portal release (Release 28 (26-11-2019)). Patients with sufficient clinical data that received gemcitabine-based adjuvant therapy were included. All patients included underwent primary surgical resection for PDAC and were defined as receiving gemcitabine-based adjuvant therapy if completing 3 cycles or more. RNAseq analysis was performed as previously described (31). Patients were dichotomized as high or low based on Variance Stabilizing Transformation (VST) normalized (using the DESEq2 package, v1.42.1) followed by Z-score transformed counts ranked from highest to lowest and split into equally sized group (31). Clinical variables were determined using the AJCC 7^th^ staging system. Survival was computed using Kaplan-Meier plots and the log rank test. A gene signature consisting of *Slc29a1*, *Slc29a2*, *Slc8a3*, and *Dck* were used to define a gemcitabine uptake/metabolism signature. A cox model with interaction term was run to test for predictive versus prognostic effect of *Hmgcr*.

### qPCR

RNA samples from parental and GemR KPC-mT3 and KPC-FC1242 cells were isolated using the RNeasy Plus Mini Kit (Qiagen) according to the manufacturer’s instructions. RNA purity was assessed using a NanoDrop 2000c (Thermo Scientific). RNA samples (1 µg) were reverse transcribed to cDNA using the iScript cDNA Synthesis Kit (Bio-Rad). qPCR was performed on the QuantStudio 5 PCR System with QuantStudio Design and Analysis Software v1.5.2 (Thermo Fisher Scientific) using Fast SYBR Green Master Mix (Thermo Fisher). Primer sequences are listed below. The gene expression was calculated as delta delta CT. TBP and HPRT1 were used as housekeeping genes.

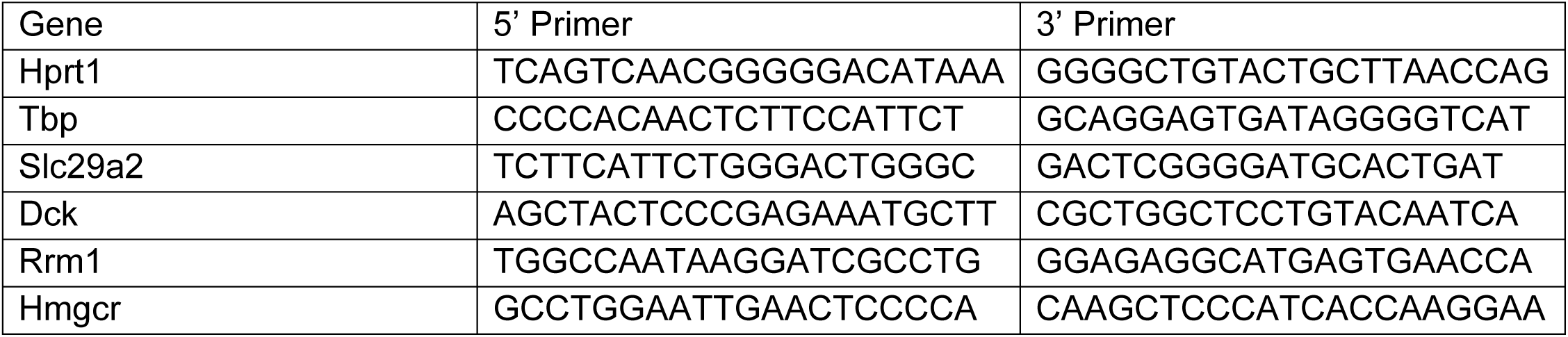
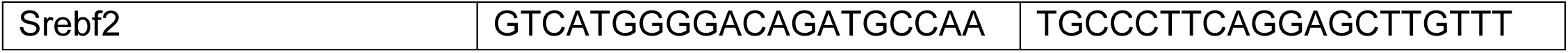

### Western blotting

All lysates were quantified via bicinchoninic acid assay (Thermo Fisher Scientific). Protein amounts were equalized and run on native PAGE gels (Bio-Rad Mini-PROTEAN), then transferred to nitrocellulose membranes (Bio-Rad Transblot) and blocked with 5% milk. After washing and incubation in primary antibodies, membranes were washed and incubated in secondary antibodies. Membranes were then visualized on a Thermo Fisher iBright CL1500 (1.7.0) using chemiluminescent substrates (Supersignal™ West Pico PLUS or Supersignal™ West Femto, Thermo Fisher Scientific). Quantification was performed using ImageJ software.

The antibodies used in this study are detailed below.

| Antibody | Vendor | Catalog Number | Application | Dilution |
| --- | --- | --- | --- | --- |
| γH2AX | Cell Signaling | 9718 | Western blot | 1:1000 |
| Cleaved Caspase 3 (CC3) | Cell Signaling | 9664 | Western blot | 1:1000 |
| Vinculin | Cell Signaling | 13901 | Western blot | 1:5000 |
| SREBF2 | Thermo Fisher | PA1-338 | Western blot | 1:1000 |
| HMGCR | Thermo Fisher | MA5-35242 | Western Blot | 1:1000 |
| β-actin | Santa Cruz | Sc-4778 | Western Blot | 1:5000 |
| HDJ2 | Novus Biologicals | NB600-952 | Western Blot | 1:1000 |
| Unprenylated RAP1a | Santa Cruz | sc-373968 | Western Blot | 1:200 |

### Subcutaneous allograft assay

Parental and GemR KPC-mT3, and parental and GemR KPC-FC1199 cells, respectively, were implanted by subcutaneous injection of 5 × 10^5^ cells in 200 µL of 1:1 serum-free DMEM:BME (Cultrex® Reduced Growth Factor Basement Membrane Matrix, Type R1, Fisher) into the flanks of 8 to 12-week-old male C57Bl/6J mice. For each treatment group, *n* = 8 allografts were implanted. Treatment was initiated on day 7 with gemcitabine (60 mg/kg dissolved in RNAse free H_2_O) or vehicle by i.p. injection every three days. Mice were euthanized on day 18 (KPC-FC1199), or day 21 (KPC-mT3), respectively, or when a predefined ethical endpoint was reached. Tumor size was measured with a caliper twice per week. Tumor volumes were determined with the following equation: *v* = (*l* × *w*^2^) × π / 6 (*v* is volume, *l* is length, and *w* is width).

Parental and GemR KPC-mT3 cells were implanted by subcutaneous injection of 1 x 10^6^ (parental KPC-mT3) and 5 × 10^5^ (GemR KPC-mT3) cells in 200 µL of 1:1 serum-free DMEM:BME (Cultrex® Reduced Growth Factor Basement Membrane Matrix, Type R1, Fisher) into the flanks of 8 to 12-week-old male C57Bl/6J mice. For each treatment group, *n* ≥ 10 allografts were implanted. Mice were fed a liquid diet (Ensure Plus, Abbott Nutrition) beginning the day of tumor cell injection. Treatment was initiated on day 7 (GemR KPC-mT3) or day 8 (parental KPC-mT3). Treatment consisted of gemcitabine injected intraperitoneally every three days (60 mg/kg dissolved in RNAse free H_2_O), pitavastatin administered by oral gavage twice daily (60 mg/kg dissolved in 0.1% Tween/0.5% methylcellulose/RNAse free H_2_O), gemcitabine plus pitavastatin, or vehicle. Mice were euthanized on day 18 or when a predefined ethical endpoint was reached. Tumor size was measured with a caliper twice per week. Tumor volumes were determined with the following equation: *v* = (*l* × *w*^2^) × π / 6 (*v* is volume, *l* is length, and *w* is width).

### Orthotopic assay

Gemcitabine-resistant KPC-mT3 cells (2 × 10^3^) were suspended in 50 µl of a 4:1 BME:DMEM (Cultrex® Reduced Growth Factor Basement Membrane Matrix, Type R1, Fisher) mixture and injected orthotopically into 8 to 12-week-old male C57BL/6J mice. For each treatment group, *n* ≥ 8 allografts were implanted. Mice were fed a liquid diet (Ensure Plus, Abbott Nutrition) beginning the day of tumor cell injection. Treatment was initiated on day 7. Treatment consistent of gemcitabine injected intraperitoneally every three days (60 mg/kg dissolved in RNAse free H_2_O), pitavastatin administered by oral gavage twice daily (60 mg/kg dissolved in 0.1% Tween/0.5% methylcellulose/RNAse free H_2_O), gemcitabine plus pitavastatin, or vehicle. Mice were euthanized on day 17, or when a predefined ethical endpoint was reached.

### Histology

After euthanization, resected tumors were fixed overnight with Z-fix solution (Anatech). Tumors were processed using a Leica TP1020 tissue processor, embedded in paraffin, and cut into 5 µm sections. Automated immunohistochemical staining was performed by the Department of Pathology and Laboratory Medicine at UC Irvine Health using a Benchmark ULTRA Stainer (Roche Diagnostics). Phospho-Histone H2A.X (Cell Signaling) was used for staining. H2AX p-S139-positive areas of bright-field images were quantified using ImageJ software (IHC image analysis toolbox, normalization of positive areas to the total surface to calculate the respective percentage occupied by positive staining per field). All quantifications were performed on 5 arbitrary images of 4 subcutaneous tumors included in the study.

### Data availability

RNA sequencing data will be uploaded to GEO upon acceptance of manuscript.

### Statistical analysis

GraphPad Prism software was used for statistical analysis. Groups of two were analyzed with two-tailed Student’s t-test, groups greater than two with a single variable were compared using one-way ANOVA analysis with Tukey post hoc test, and groups greater than two with multiple variables were compared with two-way ANOVA with Tukey post hoc test. The following values were used to denote statistical significance: *, *P* ≤ 0.05; **, *P* ≤ 0.01; ***, *P* ≤ 0.001; and ****, *P* ≤ 0.0001.

## Acknowledgments

We would like to thank Devon Pendlebury, Sebastian Michels, Johann Gout, Elodie Roger and the members of the Halbrook Lab for support and critical reading of the manuscript. We are grateful to Andrea Wißmann for her excellent technical support, and to Jessica Lindenmayer, Tapan Joshi and Verena Renz from the Core Facility Organoids of the Medical Faculty at Ulm University for their outstanding technical assistance. C.J.H. was supported by R00CA241357, R37CA283575, American Cancer Society Research Scholar Grant (RSG-1255258), a V Scholar award (V2021-026), a UC Cancer Research Coordinating Committee Faculty Seed Grant, a Sky Foundation Award, and a Collaborative Award from the University of California Pancreatic Cancer Consortium. C.J.H, C.J., A.E., T.F.M, J.B.V., and D.F were supported by the Chao Family Comprehensive Cancer Center support grant (P30CA062203). A.K.B was supported by a fellowship through the German Cancer Aid Foundation (Mildred-Scheel-Postdoktorandenstipendium). S.C was supported by T32CA009054. T.F.M. was supported by K01CA249038. A.M. was supported by the National Cancer Institute (R01 CA276461) and the V Foundation (V Scholar Award).

## Author Contributions

CJH and AKB conceived of, designed this study, and wrote the manuscript. CJH, SC, EN, KG, LPSS, EZ, RS, CA, NY, CA, GT, CG, AC, DJ, TS, AM, CJ, ALE, MS, TFM, AK, DKC, HT, JBV, and DAF provided key reagents, performed experiments, analyzed, and interpreted data. CJH supervised the work carried out in this study and obtained funding.

---

**Supplementary Figure 1.**
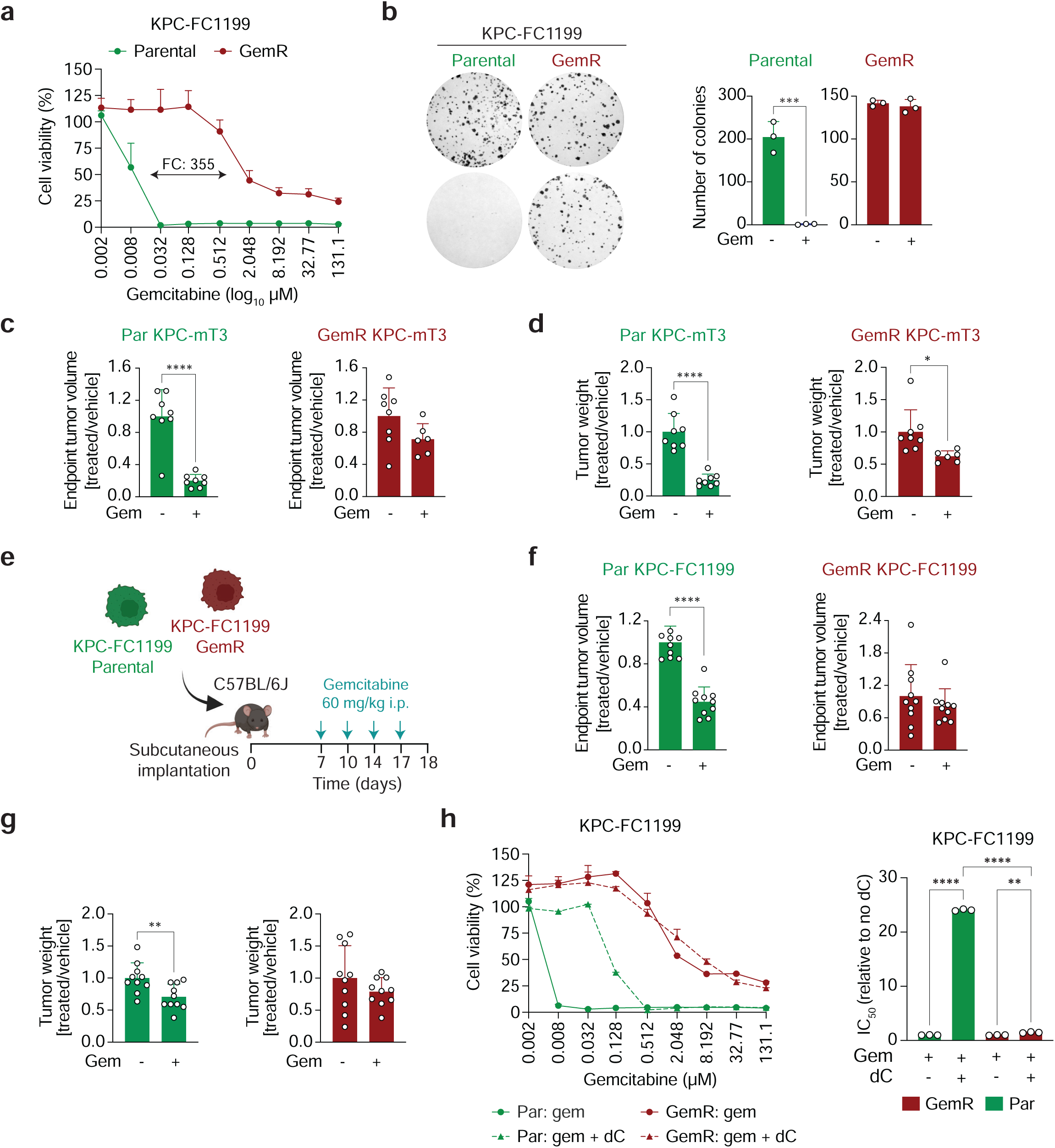
**a.** Gemcitabine dose response curves for parental (Par) and gemcitabine-resistant (GemR) KPC-FC1199 cells, with fold change (FC) indicated. **b.** Representative images and quantification of colony formation assays of vehicle or 0.01 µM gemcitabine (gem) treatment in parental and GemR KPC-FC1299 cells. **c-d.** Endpoint tumor volume (**c**) and tumor weight (**d**) of Par and GemR KPC-mT3 allografts from subcutaneous experiment shown in (Figure 1g). **e.** Treatment schematic of subcutaneous in vivo experiment with parental and GemR KPC-FC1199 cells. **f-g.** Endpoint tumor volume (**f**) and tumor weight (**g**) of Par and GemR KPC-FC1199 allografts from subcutaneous experiment shown in (**Supplementary Figure 1e**). **h.** Dose response curves and quantification for parental and GemR KPC-FC1199 cells treated with gemcitabine +/- 10 µM deoxycytidine (dC). Error bars are presented as mean ± SD (**b, c, d, f, g**). * *P* ≤ 0.05; ** *P* ≤ 0.01; *** *P* ≤ 0.001; **** *P* ≤ 0.0001 by two-tailed Student’s t-test (**c, d, f, g**).

**Supplementary Figure 2.**
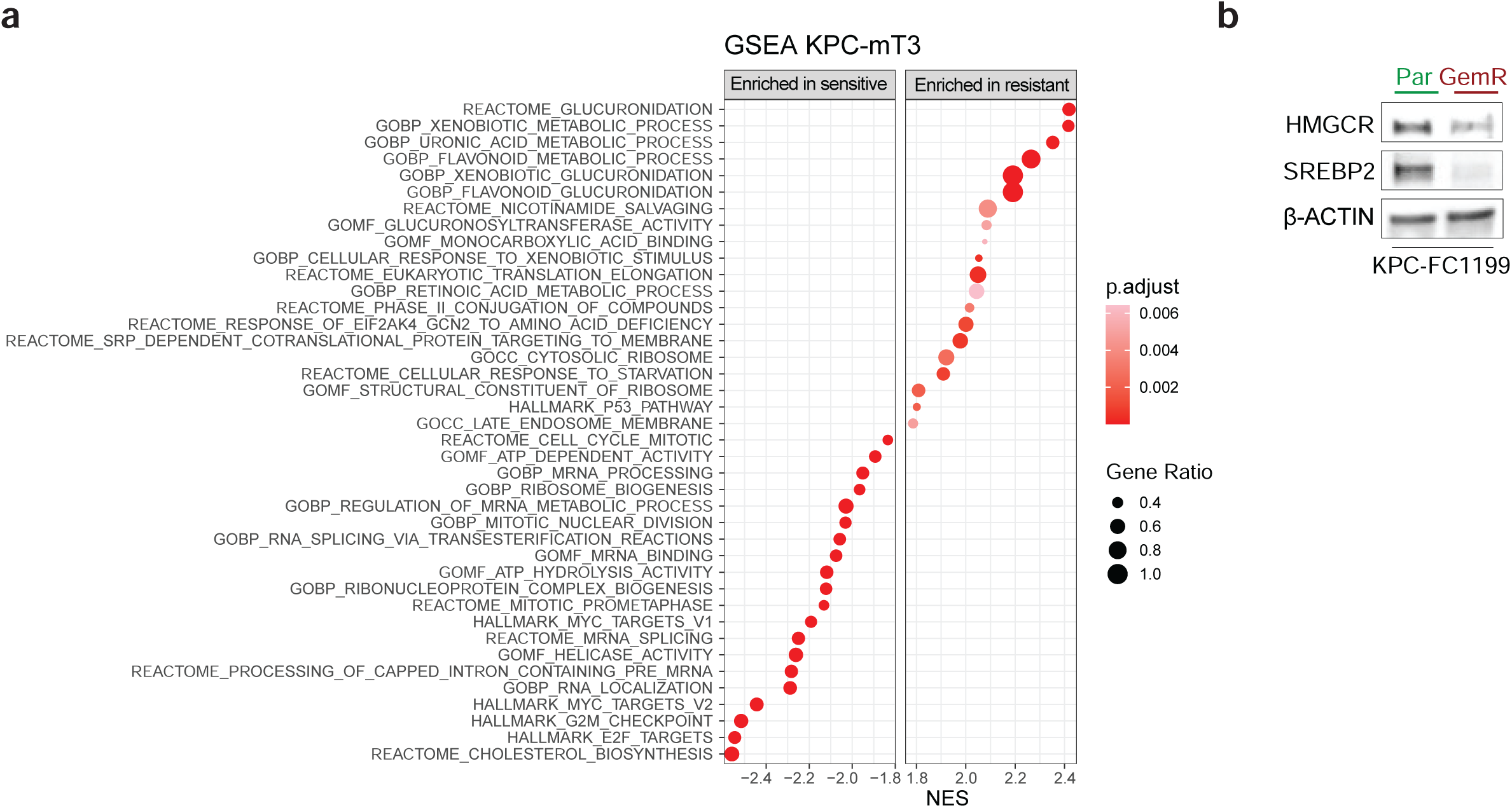
**a.** Gene set enrichment analysis in gemcitabine-resistant (GemR) KPC-mT3 cells. **b.** Representative immunoblot for HMGCR and SREBP2 on parental (Par) and GemR KPC-FC1199 cell lysates.

**Supplementary Figure 3.**
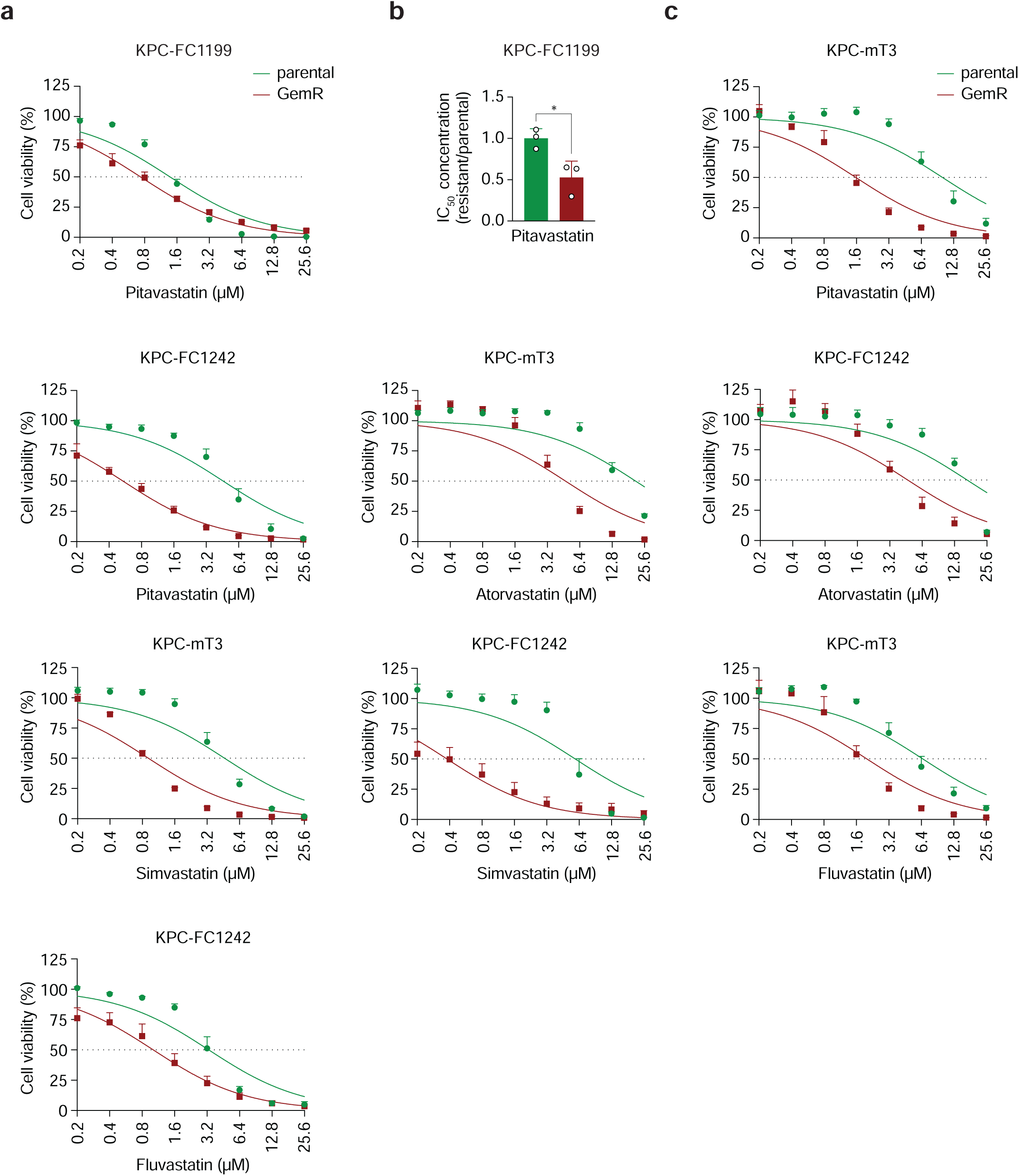
**a.** IC_50_ curves for pitavastatin in parental (Par) and gemcitabine-resistant (GemR) KPC-FC1199 cells. **b.** IC_50_ concentration for pitavastatin normalized to Par KPC-FC1199 cells. **c.** IC_50_ curves for pitavastatin, atorvastatin, simvastatin, and fluvastatin in parental and GemR KPC-mT3 and KPC-FC1242 cells. Error bars are presented as mean ± SEM (**a, c**) or mean ± SD (**b**). * *P* ≤ 0.05 by two-tailed Student’s t-test (**b**).

**Supplementary Figure 4.**
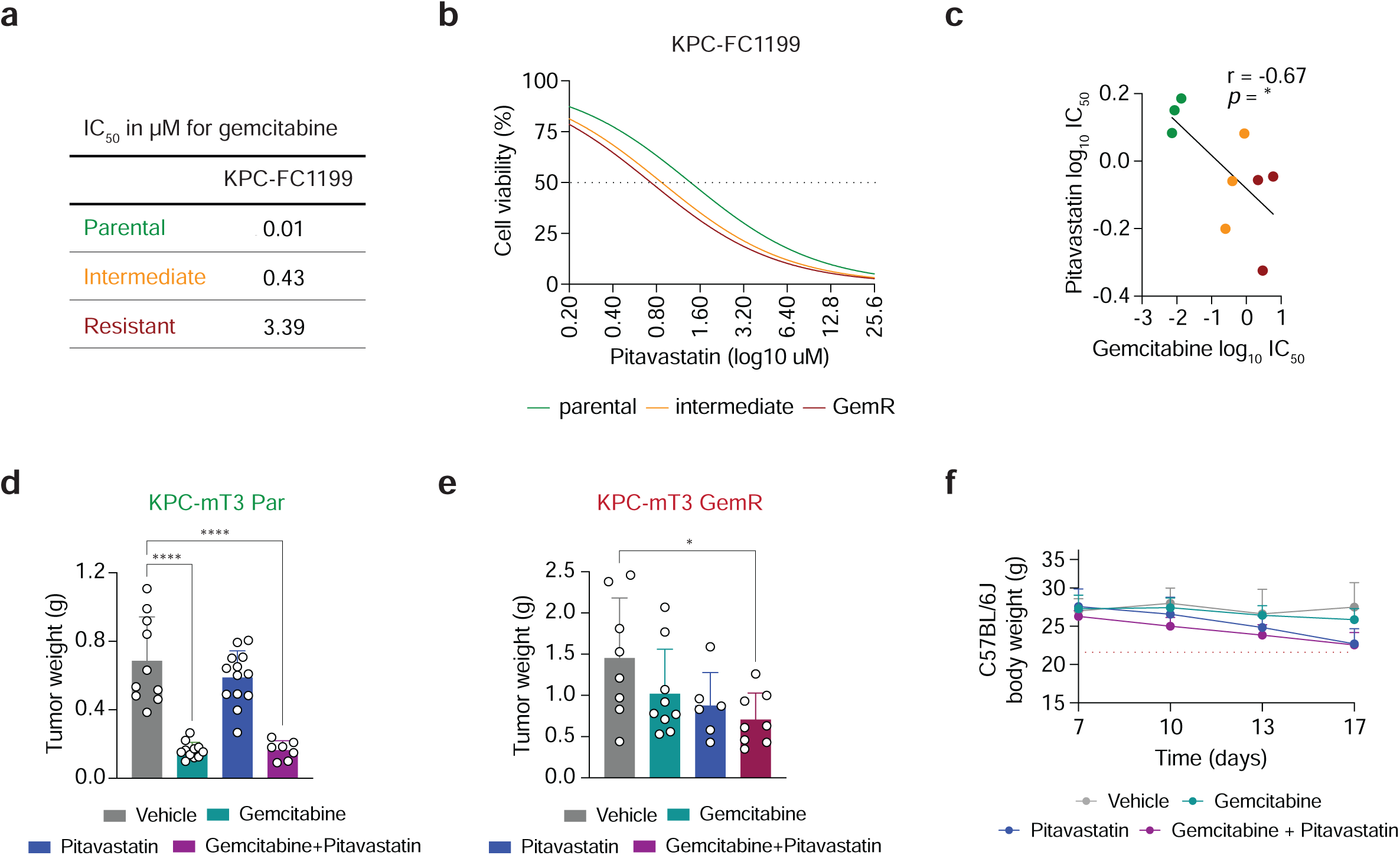
**a.** Table with IC_50_ concentrations for gemcitabine in parental, intermediate resistant, and GemR KPC-FC1199 cells. **b-c**. IC_50_ curves for pitavastatin (**b**) and correlation of IC_50_ concentrations for pitavastatin and gemcitabine (**c**) of parental, intermediate resistant, and GemR KPC-FC1199 cells. **d**. Tumor weight of parental (Par) KPC-mT3 (**d**) and GemR KPC-mT3 (**e**) allografts from subcutaneous experiment shown in (Figure 4i). **f.** C57BL/6J mouse weight under treatments from orthotopic experiment shown in (Figure 4l). Error bars are presented as mean ± SD (**d, e, f**). * *P* ≤ 0.05; *** *P* ≤ 0.001; **** *P* ≤ 0.0001 by Pearson correlation (**c**) and one-way ANOVA with Tukey post hoc test (**d, e**).

**Supplementary Figure 5.**
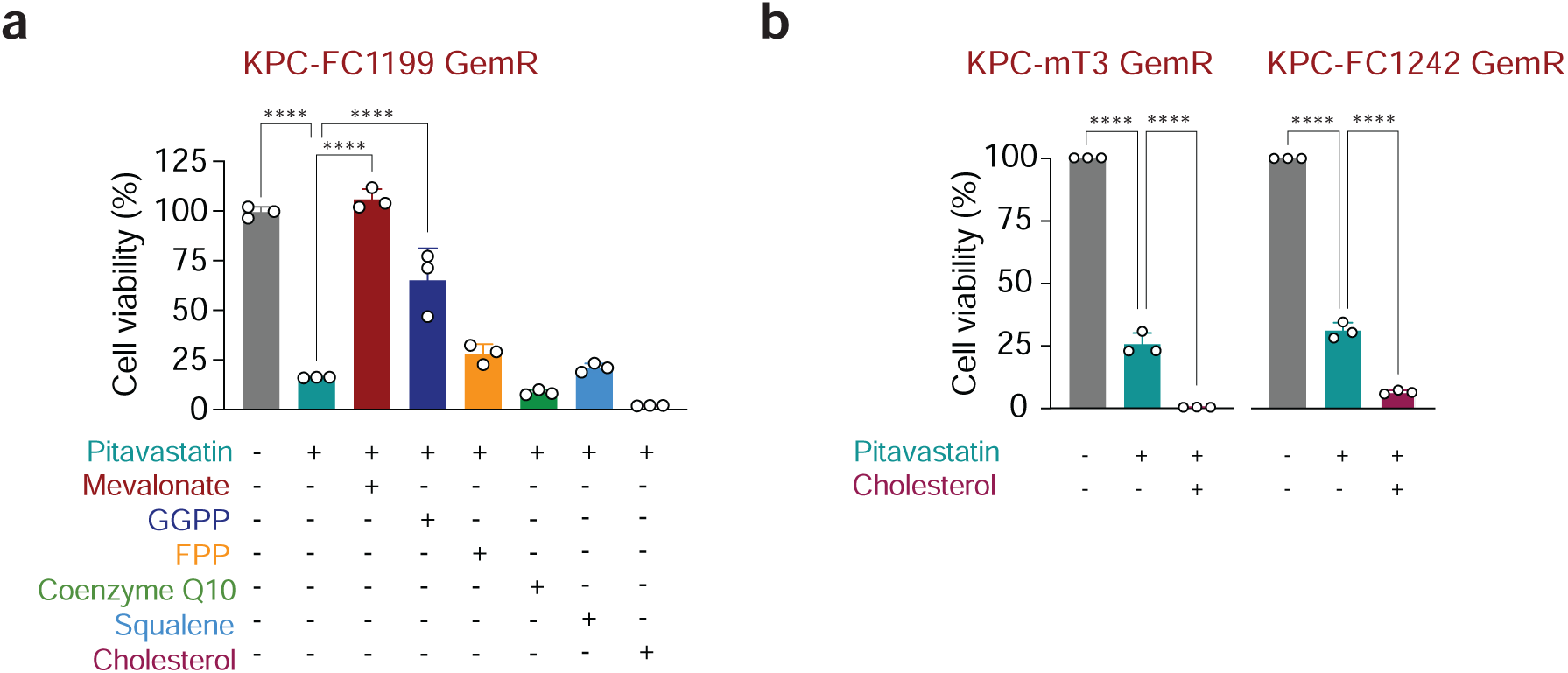
**a.** Cell viabilities of gemcitabine-resistant (GemR) KPC-FC1199 cells treated with 3 µM pitavastatin alone or in combination with 1 mM mevalonate, 20 µM GGPP, 20 µM FPP, 10 µM coenzyme Q10, 50 µM squalene, or 50 µM cholesterol. **b.** Cell viabilities of GemR KPC-mT3 and GemR KPC-FC1242 cells treated with 5 µM pitavastatin alone or in combination 50 µM cholesterol. Error bars are presented as mean ± SD (**a, b**). **** *P* ≤ 0.0001 by one-way ANOVA with Tukey post hoc test (**a,b**).

## References

1. Siegel RL, Kratzer TB, Giaquinto AN, Sung H, Jemal A. Cancer statistics, 2025. CA Cancer J Clin. 2025;75(1):10–45.

2. Halbrook CJ, Lyssiotis CA, Pasca di Magliano M, Maitra A. Pancreatic cancer: Advances and challenges. Cell. 2023;186(8):1729–54.

3. Von Hoff DD, Ervin T, Arena FP, Chiorean EG, Infante J, Moore M, et al. Increased survival in pancreatic cancer with nab-paclitaxel plus gemcitabine. N Engl J Med. 2013;369(18):1691–703.

4. Conroy T, Desseigne F, Ychou M, Bouche O, Guimbaud R, Becouarn Y, et al. FOLFIRINOX versus gemcitabine for metastatic pancreatic cancer. N Engl J Med. 2011;364(19):1817–25.

5. Wainberg ZA, Melisi D, Macarulla T, Pazo Cid R, Chandana SR, De La Fouchardiere C, et al. NALIRIFOX versus nab-paclitaxel and gemcitabine in treatment-naive patients with metastatic pancreatic ductal adenocarcinoma (NAPOLI 3): a randomised, open-label, phase 3 trial. Lancet. 2023;402(10409):1272–81.

6. Kamphorst JJ, Nofal M, Commisso C, Hackett SR, Lu W, Grabocka E, et al. Human pancreatic cancer tumors are nutrient poor and tumor cells actively scavenge extracellular protein. Cancer Res. 2015;75(3):544–53.

7. Provenzano PP, Cuevas C, Chang AE, Goel VK, Von Hoff DD, Hingorani SR. Enzymatic Targeting of the Stroma Ablates Physical Barriers to Treatment of Pancreatic Ductal Adenocarcinoma. Cancer Cell. 2012;21(3):418–29.

8. Sullivan MR, Danai LV, Lewis CA, Chan SH, Gui DY, Kunchok T, et al. Quantification of microenvironmental metabolites in murine cancers reveals determinants of tumor nutrient availability. Elife. 2019;8.

9. Halbrook CJ, Lyssiotis CA. Employing Metabolism to Improve the Diagnosis and Treatment of Pancreatic Cancer. Cancer Cell. 2017;31(1):5–19.

10. Perera RM, Bardeesy N. Pancreatic Cancer Metabolism: Breaking It Down to Build It Back Up. Cancer Discov. 2015;5(12):1247–61.

11. Kim PK, Halbrook CJ, Kerk SA, Radyk M, Wisner S, Kremer DM, et al. Hyaluronic acid fuels pancreatic cancer cell growth. Elife. 2021;10.

12. Sousa CM, Biancur DE, Wang X, Halbrook CJ, Sherman MH, Zhang L, et al. Pancreatic stellate cells support tumour metabolism through autophagic alanine secretion. Nature. 2016;536(7617):479–83.

13. Banh RS, Biancur DE, Yamamoto K, Sohn ASW, Walters B, Kuljanin M, et al. Neurons Release Serine to Support mRNA Translation in Pancreatic Cancer. Cell. 2020;183(5):1202–18 e25.

14. Parker SJ, Amendola CR, Hollinshead KER, Yu Q, Yamamoto K, Encarnacion-Rosado J, et al. Selective Alanine Transporter Utilization Creates a Targetable Metabolic Niche in Pancreatic Cancer. Cancer Discov. 2020;10(7):1018–37.

15. Halbrook CJ, Thurston G, Boyer S, Anaraki C, Jimenez JA, McCarthy A, et al. Differential integrated stress response and asparagine production drive symbiosis and therapy resistance of pancreatic adenocarcinoma cells. Nat Cancer. 2022;3(11):1386–403.

16. Halbrook CJ, Pontious C, Kovalenko I, Lapienyte L, Dreyer S, Lee HJ, et al. Macrophage-Released Pyrimidines Inhibit Gemcitabine Therapy in Pancreatic Cancer. Cell Metab. 2019;29(6):1390–9 e6.

17. Beutel AK, Halbrook CJ. Barriers and opportunities for gemcitabine in pancreatic cancer therapy. Am J Physiol Cell Physiol. 2023;324(2):C540–C52.

18. Dalin S, Sullivan MR, Lau AN, Grauman-Boss B, Mueller HS, Kreidl E, et al. Deoxycytidine Release from Pancreatic Stellate Cells Promotes Gemcitabine Resistance. Cancer Res. 2019;79(22):5723–33.

19. Shukla SK, Purohit V, Mehla K, Gunda V, Chaika NV, Vernucci E, et al. MUC1 and HIF-1alpha Signaling Crosstalk Induces Anabolic Glucose Metabolism to Impart Gemcitabine Resistance to Pancreatic Cancer. Cancer Cell. 2017;32(3):392.

20. Son J, Lyssiotis CA, Ying H, Wang X, Hua S, Ligorio M, et al. Glutamine supports pancreatic cancer growth through a KRAS-regulated metabolic pathway. Nature. 2013;496(7443):101-5.

21. Juarez D, Fruman DA. Targeting the Mevalonate Pathway in Cancer. Trends Cancer. 2021;7(6):525–40.

22. Lee JS, Roberts A, Juarez D, Vo TT, Bhatt S, Herzog LO, et al. Statins enhance efficacy of venetoclax in blood cancers. Sci Transl Med. 2018;10(445).

23. Hillis AL, Martin TD, Manchester HE, Hogstrom J, Zhang N, Lecky E, et al. Targeting Cholesterol Biosynthesis with Statins Synergizes with AKT Inhibitors in Triple-Negative Breast Cancer. Cancer Res. 2024;84(19):3250–66.

24. Guo C, Wan R, He Y, Lin SH, Cao J, Qiu Y, et al. Therapeutic targeting of the mevalonate-geranylgeranyl diphosphate pathway with statins overcomes chemotherapy resistance in small cell lung cancer. Nat Cancer. 2022;3(5):614–28.

25. Brem EA, O’Brien SM, Becker PS, Jeyakumar D, Pinter-Brown LC, Kirk C, et al. A Phase 1 Study of Adding Pitavastatin to Venetoclax-Based Therapy in AML and CLL/SLL. Blood. 2023;142.

26. Li Y, Tang S, Wang H, Zhu H, Lu Y, Zhang Y, et al. A pancreatic cancer organoid biobank links multi-omics signatures to therapeutic response and clinical evaluation of statin combination therapy. Cell Stem Cell.

27. Abdullah MI, de Wolf E, Jawad MJ, Richardson A. The poor design of clinical trials of statins in oncology may explain their failure - Lessons for drug repurposing. Cancer Treat Rev. 2018;69:84–9.

28. Schachter M. Chemical, pharmacokinetic and pharmacodynamic properties of statins: an update. Fundam Clin Pharmacol. 2005;19(1):117–25.

29. de Wolf E, Abdullah MI, Jones SM, Menezes K, Moss DM, Drijfhout FP, et al. Dietary geranylgeraniol can limit the activity of pitavastatin as a potential treatment for drug-resistant ovarian cancer. Sci Rep. 2017;7(1):5410.

30. Goldstein JL, Brown MS. Regulation of the mevalonate pathway. Nature. 1990;343(6257):425–30.

31. Bailey P, Chang DK, Nones K, Johns AL, Patch AM, Gingras MC, et al. Genomic analyses identify molecular subtypes of pancreatic cancer. Nature. 2016;531(7592):47–52.

32. Kubota CS, Myers SL, Seppala TT, Burkhart RA, Espenshade PJ. In vivo CRISPR screening identifies geranylgeranyl diphosphate as a pancreatic cancer tumor growth dependency. Mol Metab. 2024;85:101964.

33. Berndt N, Hamilton AD, Sebti SM. Targeting protein prenylation for cancer therapy. Nat Rev Cancer. 2011;11(11):775–91.

34. Dudakovic A, Wiemer AJ, Lamb KM, Vonnahme LA, Dietz SE, Hohl RJ. Inhibition of geranylgeranyl diphosphate synthase induces apoptosis through multiple mechanisms and displays synergy with inhibition of other isoprenoid biosynthetic enzymes. J Pharmacol Exp Ther. 2008;324(3):1028–36.

35. Gabitova-Cornell L, Surumbayeva A, Peri S, Franco-Barraza J, Restifo D, Weitz N, et al. Cholesterol Pathway Inhibition Induces TGF-β Signaling to Promote Basal Differentiation in Pancreatic Cancer. Cancer Cell. 2020;38(4):567–83.e11.

36. Paul JK, Azmal M, Talukder OF, Ghosh A. Statin use and Pancreatic Cancer: A Meta-analysis of its Association with Incidence in the General Population and Survival in Patients. J Gastrointest Cancer. 2025;56(1):121.

37. Tamburrino D, Crippa S, Partelli S, Archibugi L, Arcidiacono PG, Falconi M, et al. Statin use improves survival in patients with pancreatic ductal adenocarcinoma: A meta-analysis. Dig Liver Dis. 2020;52(4):392–9.

38. Abdel-Rahman O. Statin treatment and outcomes of metastatic pancreatic cancer: a pooled analysis of two phase III studies. Clin Transl Oncol. 2019;21(6):810–6.

39. Lee HS, Lee SH, Lee HJ, Chung MJ, Park JY, Park SW, et al. Statin Use and Its Impact on Survival in Pancreatic Cancer Patients. Medicine (Baltimore). 2016;95(19):e3607.

40. Hong JY, Nam EM, Lee J, Park JO, Lee SC, Song SY, et al. Randomized double-blinded, placebo-controlled phase II trial of simvastatin and gemcitabine in advanced pancreatic cancer patients. Cancer Chemother Pharmacol. 2014;73(1):125–30.

41. Muraguchi T, Okamoto K, Mitake M, Ogawa H, Shidoji Y. Polished rice as natural sources of cancer-preventing geranylgeranoic acid. J Clin Biochem Nutr. 2011;49(1):8–15.

42. Karbowska E, Swieczkowski D, Gasecka A, Pruc M, Safiejko K, Ladny JR, et al. Statins and the risk of pancreatic cancer: A systematic review and meta-analysis of 2,797,186 patients. Cardiol J. 2024;31(2):243–50.

43. Archibugi L, Arcidiacono PG, Capurso G. Statin use is associated to a reduced risk of pancreatic cancer: A meta-analysis. Dig Liver Dis. 2019;51(1):28–37.

44. Ardito CM, Gruner BM, Takeuchi KK, Lubeseder-Martellato C, Teichmann N, Mazur PK, et al. EGF receptor is required for KRAS-induced pancreatic tumorigenesis. Cancer Cell. 2012;22(3):304–17.

45. Daniluk J, Liu Y, Deng D, Chu J, Huang H, Gaiser S, et al. An NF-kappaB pathway-mediated positive feedback loop amplifies Ras activity to pathological levels in mice. J Clin Invest. 2012;122(4):1519–28.

46. Park JH, Mortaja M, Son HG, Zhao X, Sloat LM, Azin M, et al. Statin prevents cancer development in chronic inflammation by blocking interleukin 33 expression. Nat Commun. 2024;15(1):4099.

47. Guerra C, Collado M, Navas C, Schuhmacher AJ, Hernandez-Porras I, Canamero M, et al. Pancreatitis-induced inflammation contributes to pancreatic cancer by inhibiting oncogene-induced senescence. Cancer Cell. 2011;19(6):728–39.

48. Ritchie ME, Phipson B, Wu D, Hu Y, Law CW, Shi W, et al. limma powers differential expression analyses for RNA-sequencing and microarray studies. Nucleic Acids Res. 2015;43(7):e47.

49. Yu G, Wang LG, Han Y, He QY. clusterProfiler: an R package for comparing biological themes among gene clusters. OMICS. 2012;16(5):284–7.

50. Ge SX, Jung D, Yao R. ShinyGO: a graphical gene-set enrichment tool for animals and plants. Bioinformatics. 2019;36(8):2628–9.

51. Dreyer SB, Jamieson NB, Upstill-Goddard R, Bailey PJ, McKay CJ, Australian Pancreatic Cancer Genome I, et al. Defining the molecular pathology of pancreatic body and tail adenocarcinoma. Br J Surg. 2018;105(2):e183–e91.

